# ComM is a hexameric helicase that promotes branch migration during natural transformation in diverse Gram-negative species

**DOI:** 10.1101/147660

**Authors:** Thomas M. Nero, Triana N. Dalia, Joseph Che-Yen Wang, David T. Kysela, Matthew L. Bochman, Ankur B. Dalia

## Abstract

Acquisition of foreign DNA by natural transformation is an important mechanism of adaptation and evolution in diverse microbial species. Here, we characterize the mechanism of ComM, a broadly conserved AAA+ protein previously implicated in homologous recombination of transforming DNA (tDNA) in naturally competent Gram-negative bacterial species. *In vivo*, we found that ComM was required for efficient comigration of linked genetic markers in *Vibrio cholerae* and *Acinetobacter baylyi*, which is consistent with a role in branch migration. Also, ComM was particularly important for integration of tDNA with increased sequence heterology, suggesting that its activity promotes the acquisition of novel DNA sequences. *In vitro*, we showed that purified ComM binds ssDNA, oligomerizes into a hexameric ring, and has bidirectional helicase and branch migration activity. Based on these data, we propose a model for tDNA integration during natural transformation. This study provides mechanistic insight into the enigmatic steps involved in tDNA integration and uncovers the function of a protein required for this conserved mechanism of horizontal gene transfer.

## INTRODUCTION

Natural competence is a physiological state in which some bacterial species can take up free DNA from the environment. Some competent species regulate the genes required for this process and, depending on the organism, competence can be induced in response to the availability of certain nutrients, quorum sensing pathways, or by DNA damage / stress (1). The Gram-negative bacterium *Vibrio cholerae* is activated for competence during growth on chitin, a polymer of β1,4-linked N-acetyl glucosamine (2). Chitin is the primary constituent of crustacean exoskeletons and is commonly found in the aquatic environment where this facultative pathogen resides. Soluble chitin oligosaccharides indirectly induce expression of TfoX, the master regulator of competence (3,4), which regulates expression of competence-related genes in concert with HapR, the master regulator of quorum sensing (5,6). *Acinetobacter baylyi* ADP1, on the other hand, is a naturally competent Gram-negative microbe that is constitutively active for competence throughout exponential growth (7).

During competence, dsDNA is bound extracellularly, however, only a single strand of this DNA is transported into the cytoplasm. Competent bacterial species may use this ingested DNA as a source of nutrients, however, if this DNA has sufficient homology to the host chromosome, the incoming DNA can also be integrated into the bacterial genome by homologous recombination (8). This process of DNA uptake and integration is referred to as natural transformation. As a result, natural transformation is an important mechanism of horizontal gene transfer and can lead to the repair of damaged DNA or facilitate acquisition of novel genetic information. Homologous recombination of single-stranded transforming DNA (tDNA) with the host chromosome requires the function of RecA, which facilitates homology searching and initiates strand invasion of tDNA through the formation of a displacement loop (D-loop). Following RecA mediated strand invasion, DNA junctions of this D-loop can then be moved in a process known as branch migration to increase or decrease the amount of tDNA integrated. Then, by a presently unresolved mechanism, this intermediate is resolved to stably integrate tDNA into the host chromosome. The molecular details involved in the integration of tDNA downstream of RecA strand invasion, however, are poorly understood.

One previously studied gene from the competent species *Haemophilus influenzae*, *comM*, is not required for DNA uptake but is required for the integration of tDNA into the host chromosome, a phenotype consistent with ComM playing a role in homologous recombination during natural transformation (9). The function of ComM, however, has remained unclear. Here, through both *in vivo* and *in vitro* characterization of ComM in *V. cholerae* and *A. baylyi*, we uncover that this protein functions as a hexameric helicase to aid in branch migration during this conserved mechanism of horizontal gene transfer.

## MATERIALS AND METHODS

### Bacterial strains and growth conditions

The parent *V. cholerae* strain used throughout this study is E7946 (10), while the *A. baylyi* strain used is ADP1 (also known as BD413) (11). A description of all strains used in this study are listed in Table S1. Strains were routinely grown in LB broth and plated on LB agar. When required, media was supplemented with 200 µg/mL spectinomycin, 50 μg/mL kanamycin, 100 μg/mL carbenicillin, 10 µg/mL trimethoprim, or 40 μg/mL X-Gal.

### Construction of mutants and transforming DNA

Linear PCR product was constructed using splicing-by-overlap extension PCR exactly as previously described (12). All primers used to construct and detect mutant alleles are described in **Table S2**. The pBAD18 Kan plasmid used as tDNA was purified from TG1, a *recA*+ *E. coli* host. Mutants were made by cotransformation exactly as previously described (13).

### Transformation assays

Transformation assays of *V. cholerae* on chitin were performed exactly as previously described (3). Briefly, ~10^8^ CFU of mid-log *V. cholerae* were incubated statically in instant ocean (IO) medium (7g/L; aquarium systems) containing chitin from shrimp shells (Sigma) for 16-24 hours at 30°C. Then, tDNA was added (500 ng for linear products containing an antibiotic resistance cassette inserted at the non-essential locus VC1807 and 1500 ng for plasmid DNA), and reactions were incubated at 30°C for 5-16 hours. Reactions were outgrown with the addition of LB medium to each reaction by shaking at 37°C for 1-3 hours and then plated for quantitative culture onto medium to select for the tDNA (transformants) or onto plain LB (total viable counts). Transformation efficiency is shown as the number of transformants / total viable counts. In cases where no colonies were observed, efficiencies were denoted as below the limit of detection (LOD) for the assay.

Chitin-independent transformation assays were performed exactly as previously described using strains that contain an IPTG inducible P_tac_ promoter upstream of the native TfoX gene (14). Briefly, strains were grown overnight with 100 μM IPTG. Then, ~10^8^ cells were diluted into instant ocean medium, and tDNA was added. Reactions were incubated statically for 5 hours and then outgrown by adding LB and shaking at 37°C for 1-3 hours. Reactions were plated for quantitative culture as described above.

For transformation of *A. baylyi* ADP1, strains were grown overnight in LB media. Then, ~10^8^ cells were diluted into fresh LB medium, and tDNA was added (~100ng). Reactions were incubated at 30°C with agitation for 5 hours and then plated for quantitative culture as described above.

### Protein expression and purification

ComM and Pif1 were cloned into StrepII expression vectors, expressed in Rosetta 2(DE3) pLysS cells using autoinduction medium (15), and purified using Strep-Tactin Sepharose (IBA). ComM protein preparations lacked detectable nuclease activity. For details please see **Supplementary Methods**.

### Electrophoretic mobility shift assay

The ssDNA probe (BBC742) and dsDNA probe (annealed BBC742 and BBC743) were labeled using Cy5 dCTP (GE Healthcare) and Terminal deoxynucleotidyl Transferase (TdT; Promega). Reactions were composed of 10 mM Tris-HCl pH 7.5, 20 mM KCl, 1 mM DTT, 10% Glycerol, 100 *γ*g/ml BSA, 9 nM Cy5 labeled probe, purified ComM protein at the indicated concentration, and 0.1 mM ATP where indicated. Reactions were incubated at RT for 30 minutes and run on 8% Tris-borate acrylamide gel at 150 V for 45 min. Probes were detected using a Chemidoc MP imaging system (Biorad).

### Blue native PAGE

Blue native PAGE was performed essentially as previously described (16). Purified ComM (2.5 μM) was incubated for 30 min at room temperature in reaction buffer [10 mM Tris-HCl pH 7.5, 20 mM KCl, 1 mM DTT, 10% Glycerol] with 5 mM ATP and/or 5 *μ*M ssDNA (oligo ABD363) where indicated. For additional details see **Supplementary Methods**.

### Negative stain electron microscopy

For negative stain electron microscopy, sample was prepared by applying 4 μL of ComM-ATP-ssDNA solution onto a glow-discharged continuous carbon film coated copper grid (EMS) and stained with 0.75% (w/v) uranyl formate. EM micrographs were acquired using a 300 kV JEM-3200FS electron microscopy with 20-eV energy slit under low dose conditions (≤ 20 e^-^/Å^2^) on a Gatan UltraScan 4000 4k x 4k CCD camera. Additional details for EM image collection and analysis are provided in the **Supplementary Methods**. The EM electron density map has been deposited to EMDataBank.org with the accession number EMD-8575.

### Phylogenetic Trees

ComM homologs were identified using a protein BLAST search of diverse bacterial genomes followed by phylogenetic analysis. Starting with a broadly representative set of genomes adapted from Wu and Eisen (17), an initial ComM candidate pool was generated from a comprehensive protein BLAST search of all predicted CDS translations against the *V. cholera*e ComM allele with a 0.001 e-value cutoff. Subsequent sequence alignment with MUSCLE (18) and maximum likelihood phylogenetic reconstruction with FastTree (19) identified a single clade of alleles with high sequence similarity (generally well above 50%) to *V. cholerae* ComM (data not shown), with the remaining alleles excluded from further analysis. After manually pruning divergent alleles with alignments covering <70% of ComM, the retained sequences comprised the set of true ComM orthologs used in the final sequence alignment and ComM phylogeny reconstruction. For whole genome phylogenetic reconstruction, Phylosift (20) identified and aligned a default set of 36 highly conserved marker genes. FastTree was used for initial reconstruction, whereas RAxML (21) subsequently estimated the maximum likelihood phylogeny for a reduced set of representative genomes under the LG (22) substitution model with gamma-distributed rate variation.

### Helicase and branch migration assays

Helicase assay substrates were 5’ end-labeled with T4 polynucleotide kinase (T4 PNK; NEB) and γ[^32^P]-ATP. The 2-strand forked helicase substrates was annealed by combining equimolar amounts of a labeled oligonucleotide with the unlabeled complement and incubating at 37°C overnight.

The short 3-strand branch migration substrates were constructed in a 2-step annealing process. First, we incubated equimolar amounts of a labeled strand and unlabeled partial complement at 95°C for 5 min. and then slow cooled to 25°C to create a forked substrate. An equimolar amount of a third strand was then added to the resulting forked product and incubated overnight at 37°C. Following annealing, the three-stranded products were PAGE purified. Primers used for all probes and additional details can be found in the **Supplementary Methods**. Helicase and branch migration activity was then assessed by incubating the indicated concentrations of ComM, Pif1, or Hrq1 with 5 mM ATP and 0.1 nM DNA substrate in resuspension buffer (25 mM Na-HEPES (pH 7.5), 5% (v/v) glycerol, 300 mM NaOAc, 5 mM MgOAc, and 0.05% Tween-20). Reactions were incubated at 37°C for 30 min and stopped with the addition of 1x Stop-Load dye (5% glycerol, 20 mM EDTA, 0.05% SDS, and 0.25% bromophenol blue) supplemented with 400 μg/mL SDS-Proteinase K followed by a 10-min incubation at 37°C. Reactions were then separated on 8% 19:1 acrylamide:bis-acrylamide gels in TBE buffer at 10 V/cm. Gels were then dried between layers of Whatman filter paper under vacuum at 55°C for 20 mins and then exposed to a phosphor imaging screen for 24-48 hours prior to scanning on a Typhoon 9210 Variable Mode Imager to image radiolabeled DNA probes. DNA unwinding and branch migration were quantified using ImageQuant 5.2 software.

The long 3-strand recombination intermediates were generated by RecA-mediated strand exchange between circular single-stranded φX174 virion DNA (NEB) and PstI-linearized double-stranded φX174 DNA (NEB) essentially as previously described (23). Briefly, recombination reactions were conducted in strand exchange buffer (25 mM Tris acetate pH 8.0, 1mM DTT, 1% glycerol, 10 mM magnesium acetate, 3 mM potassium glutamate, 10 U/mL creatine phosphokinase, 12.5 mM phosphocreatine, and 50 µg/mL acetylated BSA). First, 7.1 µM RecA (NEB) was incubated with 44 µM (in nucleotides) circular single-stranded φX174 virion DNA (CSS) at 37°C for 10 mins. Then, 2 mM ATP and ~0.84 µM SSB (Promega) were added to the reaction and incubated at 37°C for 8 mins. Finally, the strand exchange reactions were started by the addition of ~16.7 µM (in nucleotides) of PstI-linearized φX174 and incubated at 37°C for an additional 20 mins to allow for the generation of recombination intermediates. Reactions were then deproteinated and cleaned up using a PCR purification kit (Qiagen). To assess branch migration of recombination intermediates, the deproteinated DNA was then incubated with ComM in strand exchange buffer containing 5 mM ATP at 37°C for 10 mins. Reactions were then stopped using 1X stop load dye and separated on a 0.8% agarose gel in TAE. Gels were then stained with GelGreen (Biotium), and scanned on a Typhoon 9210 Variable Mode Imager.

### DNA damage assay

For DNA damage assays, ~10^8^ cells of a midlog *Vibrio cholerae* culture in instant ocean medium were treated with the indicated concentration of MMS or MMC for 1 hour at 30°C. To determine viable counts, reactions were plated for quantitative culture on LB agar.

### GFP-ComM western blots

The indicated strains of *V. cholerae* were grown with or without 100 μM IPTG. Cell lysates were run on 10% SDS PAGE gels and electrophoretically transferred to PVDF. This membrane was then blotted with primary rabbit anti-GFP (Invitrogen) or mouse anti-RpoA (Biolegend) antibodies and a goat anti-rabbit or anti-mouse IRDye 800CW (LI-COR) secondary as appropriate. Bands were detected using a LI-COR Odyssey classic infrared imaging system.

## RESULTS

### ComM is required for integration of tDNA during natural transformation

Previously, we performed an unbiased transposon-sequencing screen (Tn-seq) in *V. cholerae* to identify genes involved in natural transformation (3). One gene identified in that screen was VC0032, which encodes a homolog of *comM* from *H. influenzae*. ComM was previously implicated in the integration of tDNA during natural transformation in *H. influenzae* (9). To determine if this was also the case in *V. cholerae*, we performed chitin-dependent natural transformation assays using two distinct sources of tDNA. One was a linear PCR product that inserts an antibiotic resistance cassette at a non-essential locus, while the other was a replicating plasmid that lacks any homology to the host genome. We hypothesized that transformation with linear product requires both DNA uptake and chromosomal integration, while transformation with the plasmid only requires DNA uptake. To test this, we transformed a recombination deficient Δ*recA* strain of *V. cholerae*, and found that, as expected, this strain could not be transformed with linear product but could be transformed with a replicating plasmid, consistent with plasmid transformation being recombination-independent (**Fig. S1**). Additionally, plasmid transformation in this assay is dependent on natural competence, as mutants in genes required for uptake (Δ*pilA*) (5) and cytoplasmic protection (Δ*dprA*) of tDNA are not transformed (5,24,25) (**Fig. S1**). Using this assay, we find that a *comM* mutant (ΔVC0032) in *V. cholerae* displays a ~100-fold reduction for transformation with linear product, while rates of plasmid transformation were equal to the WT (**Fig. 1A**). This is consistent with ComM playing a role downstream of DNA uptake, and potentially during recombination. These assays were performed on chitin to induce the natural competence of this organism. To determine if ComM is playing a role specifically downstream of competence induction, we performed a chitin-independent transformation assay by overexpressing the competence regulator TfoX. Under these conditions, a *comM* mutant still had reduced rates of transformation when transformed with a linear PCR product (**Fig. 1B**). Cumulatively, these results are consistent with ComM playing a role in the late steps of transformation downstream of DNA uptake.

**Fig. 1.**
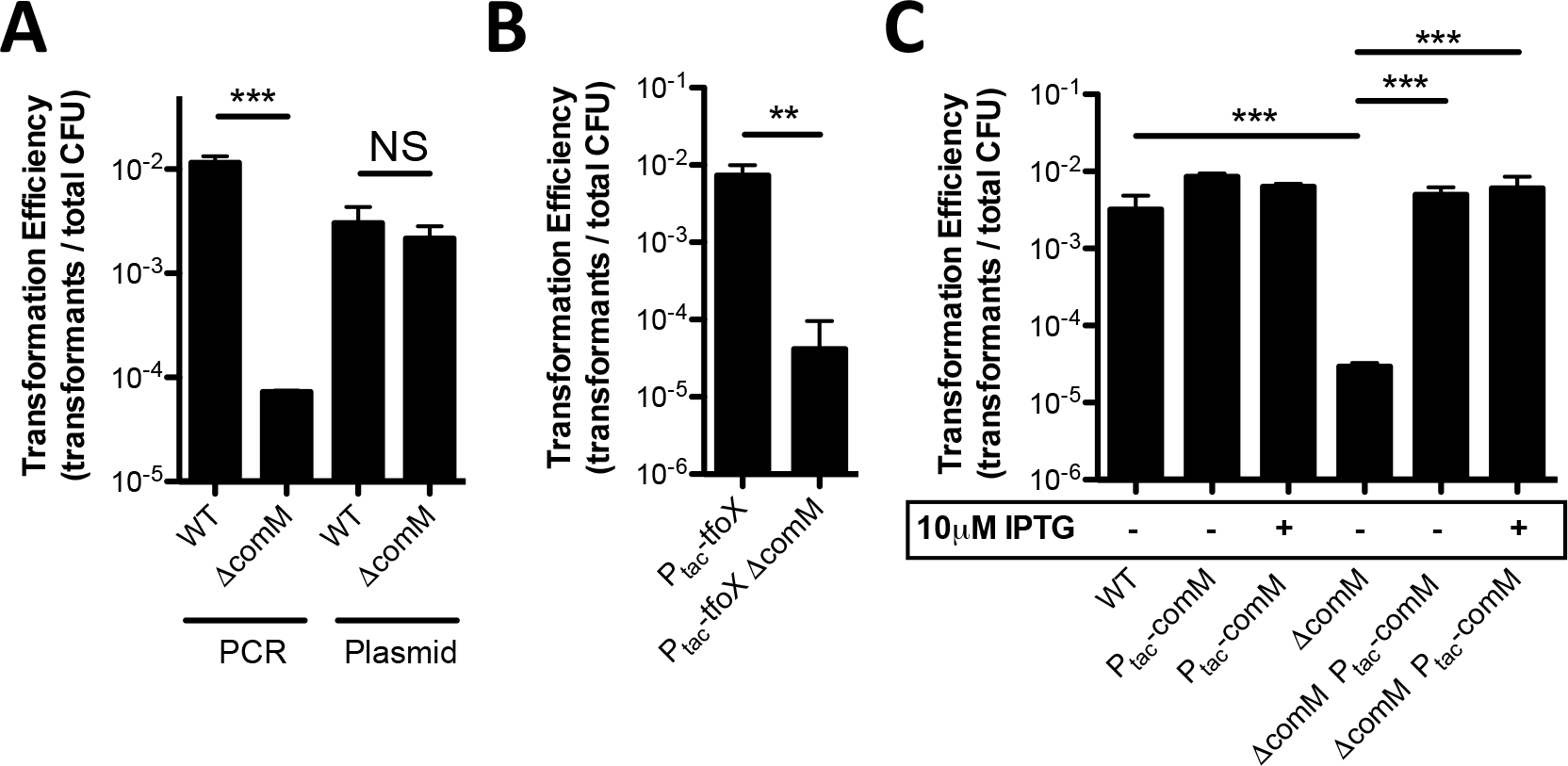
*ComM is required for integration of DNA during natural transformation*. (**A**) Chitin-dependent natural transformation assays in the indicated strains using a linear PCR product or a replicating plasmid as tDNA. (**B**) Chitin-independent transformation assays of the indicated strains with linear PCR product as tDNA. (**C**) Complementation of *comM in trans* tested in chitin-dependent transformation assays using a linear PCR product as tDNA. All data are shown as the mean ± SD and are the result of at least three independent biological replicates. ** = *p*<0.01, *** = *p*<0.001, NS = not significant

To confirm that the phenotypes observed are due to mutation of *comM*, we complemented strains by integrating an IPTG inducible P_*tac*_-*comM* construct at a heterologous site on the chromosome (at the *lacZ* locus). Our previous work has indicated that this expression construct is leaky (14) (**Fig. S2**), and consistent with this, we observe complementation even in the absence of inducer (**Fig. 1C**).

### ComM promotes branch migration through heterologous sequences in vivo

*V. cholerae* ComM is a predicted AAA+ ATPase, and members of this family have diverse functions (26). To determine if ComM had structural similarity to any AAA+ ATPase of known function, we submitted the primary sequence of this protein to the Phyre2 server (27). Despite a lack of significant homology by BLAST, this analysis revealed structural similarity to MCM2-7, the replicative helicase of eukaryotes (28,29). As a result, we explored whether ComM functions as a helicase to promote branch migration during natural transformation.

To test this *in vivo*, we assessed comigration of linked genetic markers on a linear tDNA product (**Fig. 2A**). If branch migration during transformation is efficient, we hypothesized that both markers would be integrated into the host chromosome. However, if branch migration is inefficient, we hypothesized that we may observe integration of one marker but not the other. For this assay, we generated two tDNA constructs that contained a Spec^R^marker upstream of *lacZ* as well as a genetically linked point mutation in the *lacZ* gene that was either 820bp or 245bp downstream of the Spec^R^marker (**Fig 2A**). We selected for the Spec^R^marker and screened for integration of the linked *lacZ* mutation as an indirect measure of branch migration. To prevent post-recombination repair of the *lacZ* allele by the mismatch repair (MMR) system, these experiments were performed in *mutS* mutant backgrounds. As with previous experiments, a *comM* mutant is severely reduced for transformation efficiency for the Spec^R^marker compared to the parent (**Fig. 2A, left** and **1A**). Of those cells that integrated the Spec^R^marker, the comigration efficiency of the *lacZ* mutation was higher than 90% for both products (820bp and 245bp) in the parent strain background (**Fig. 2A, right**), which is consistent with highly efficient branch migration in this background. When *comM* is deleted, however, the comigration efficiency for the *lacZ* mutation drops significantly for both products compared to the parent strain, and the reduction is more severe for the product where the *lacZ* mutation is farther away (**Fig. 2A, right**). Cumulatively, these data suggest that *comM* may play a role in branch migration during natural transformation to increase the amount of tDNA integrated into the host chromosome.

**Fig. 2.**
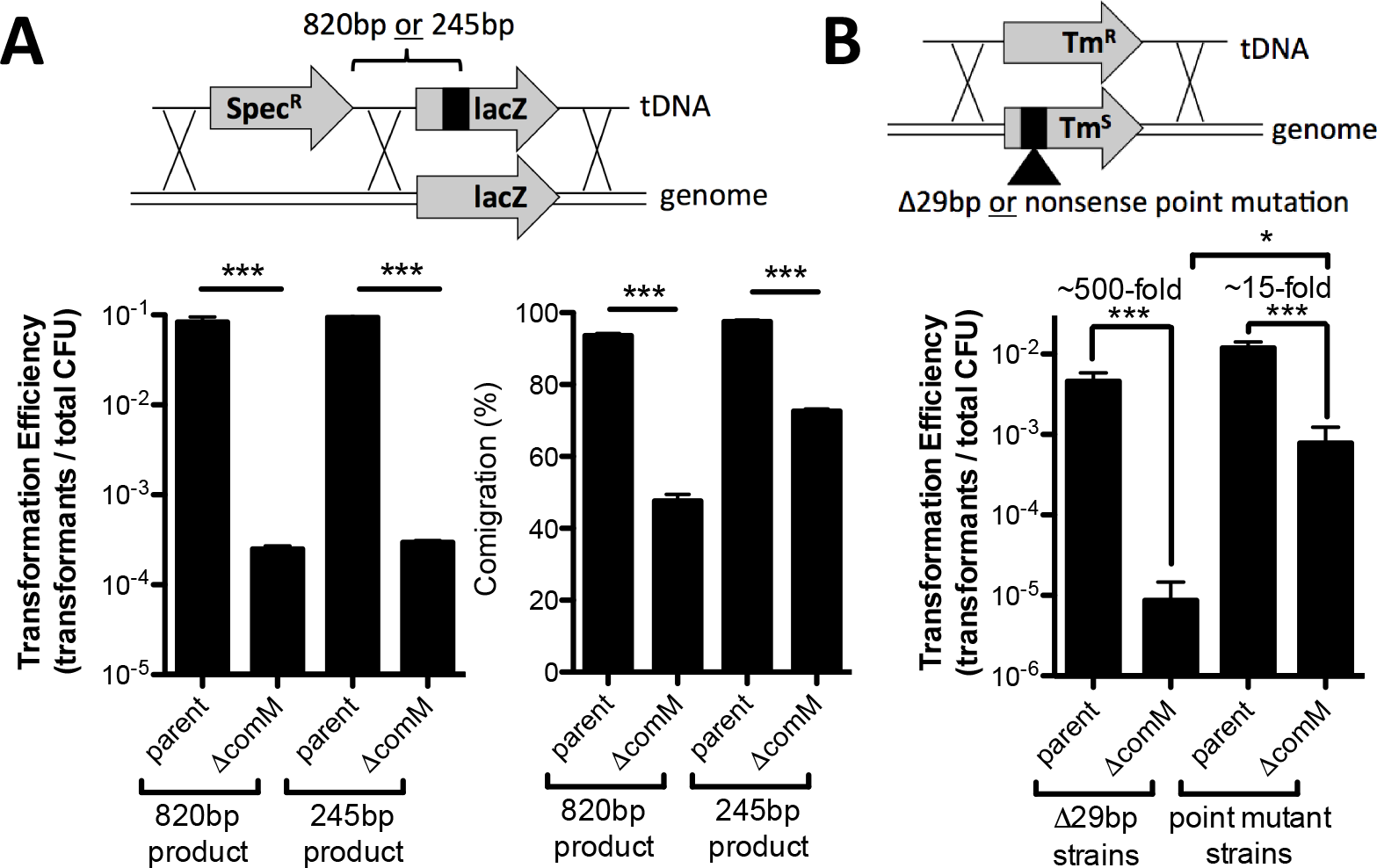
*ComM promotes branch migration through heterologous sequences* in vivo. (**A**) Chitin-dependent transformation assay performed using tDNA that contained linked genetic markers separated by 820 bp or 245 bp. (**B**) Chitin-dependent transformation assay performed in Tm^s^ strains (Tm^R^ marker inactivated by a nonsense point mutation or 29-bp deletion) using tDNA that would revert the integrated marker to Tm^R^. All strains in **A** and **B** contained a mutation in *mutS* to prevent MMR activity. In schematics above bar graphs, X’s denote possible crossover points for homologous recombination. All data are shown as the mean ± SD and are the result of at least three independent biological replicates. *** = *p*<0.001

To promote horizontal gene transfer, tDNA integrated during natural transformation must be heterologous to the host chromosome. So next, we decided to test whether ComM promotes integration of heterologous tDNA. To that end, we created strains that contain an inactivated Tm^R^marker integrated in the chromosome. The marker was inactivated with either a 29bp deletion or a nonsense point mutation. We then transformed these strains with tDNA that would restore the Tm^R^marker. Again, to eliminate any confounding effects of MMR, we performed these experiments in *mutS* mutant backgrounds. First, we find that integration of a point mutation is similar to a 29bp insertion in the parent strain background, indicating that in the presence of ComM, tDNA is efficiently integrated regardless of sequence heterology. In the *comM* mutant, however, we find that a point mutation is significantly easier to integrate compared to the 29bp insertion (**Fig. 2B**). This finding is consistent with ComM promoting branch migration through heterologous sequences during natural transformation; however, in its absence, the integration of heterologous sequences is unfavored.

### ComM hexamerizes in the presence of ATP and ssDNA

Because our *in vivo* data suggested that ComM acts as a branch migration factor, we next decided to test the biochemical activity of this protein *in vitro*. First, we determined that N-terminally tagged ComM (GFP-ComM) was functional *in vivo* while a C-terminally tagged fusion (ComM-GFP) was not (**Fig. S2**). Furthermore, recent studies indicate that *comM* is part of the competence regulon (30). Using a strain where *comM* was N-terminally tagged with GFP at the native locus, we found that *comM* protein levels are increased under competence inducing conditions (TfoX overexpression), indicating that native regulation of tagged *comM* is maintained (**Fig. S2**).

To characterize ComM *in vitro*, we expressed StrepII-ComM (N-terminal tag) in *E. coli* and purified it to homogeneity. The peak of recombinant ComM eluting from preparative gel filtration chromatography had a calculated molecular weight of 57 kDa (**Fig. S3)**. As the predicted mass of ComM is 61 kDa, this suggests that ComM exists as a monomer in solution. Many helicases oligomerize in their active state, including all known MCM family helicases (31). So, we next tested whether purified ComM oligomerizes *in vitro*. Because ComM is a predicted ATPase and may interact with DNA, we hypothesized that these factors may be required for its oligomerization and activity. To assess oligomerization, we performed blue-native PAGE (16). In this assay, ComM appears to oligomerize robustly in the presence of ATP and ssDNA, with some oligomerization also observed in the presence of ATP alone (**Fig. 3A**). This latter observation, however, may be due to a small amount of contaminating ssDNA that remains bound to ComM during purification. Defining the number of subunits in this oligomer was unreliable by blue-native PAGE due to lack of resolving power by the gel. However, the higher molecular weight species generated in the presence of ATP and ssDNA was likely larger than a dimer. Therefore, we attempted to observe ComM oligomers by negative stain transmission electron microscopy (TEM). In the absence of ligands, ComM particles were small and uniform, consistent with our gel filtration results and demonstrated the purity of our protein preparations. In the presence of ATP and ssDNA, we observed ring-like densities for ComM that upon 2-D averaging revealed that this protein forms a hexameric ring (**Fig. 3B** and **3C**). Furthermore, we generated a 3-D reconstruction from the negative stain TEM images of ComM in the presence of ATP and ssDNA (~13.8 Å resolution, see **Supplementary Methods**), which revealed that ComM forms a three-tiered barrel-like structure with a large opening on both ends (**Fig. 3D**). The pore on the bottom of this barrel is ~18Å, which can accommodate ssDNA but not dsDNA, while the 26 Å pore is able to accommodate dsDNA. The size of these pores, however, may be an underestimate due to the stain used during EM and/or ssDNA that may be bound and averaged into the 3-D construction. Regardless, these data suggest that ComM forms a hexameric ring, consistent with structures adopted by many AAA+ helicases (26). ComM also oligomerized in the presence of ADP and the non-hydrolysable ATP analog AMP-PNP, indicating that ATP is not a strict requirement for hexamer formation (**Fig. S4**).

**Fig. 3.**
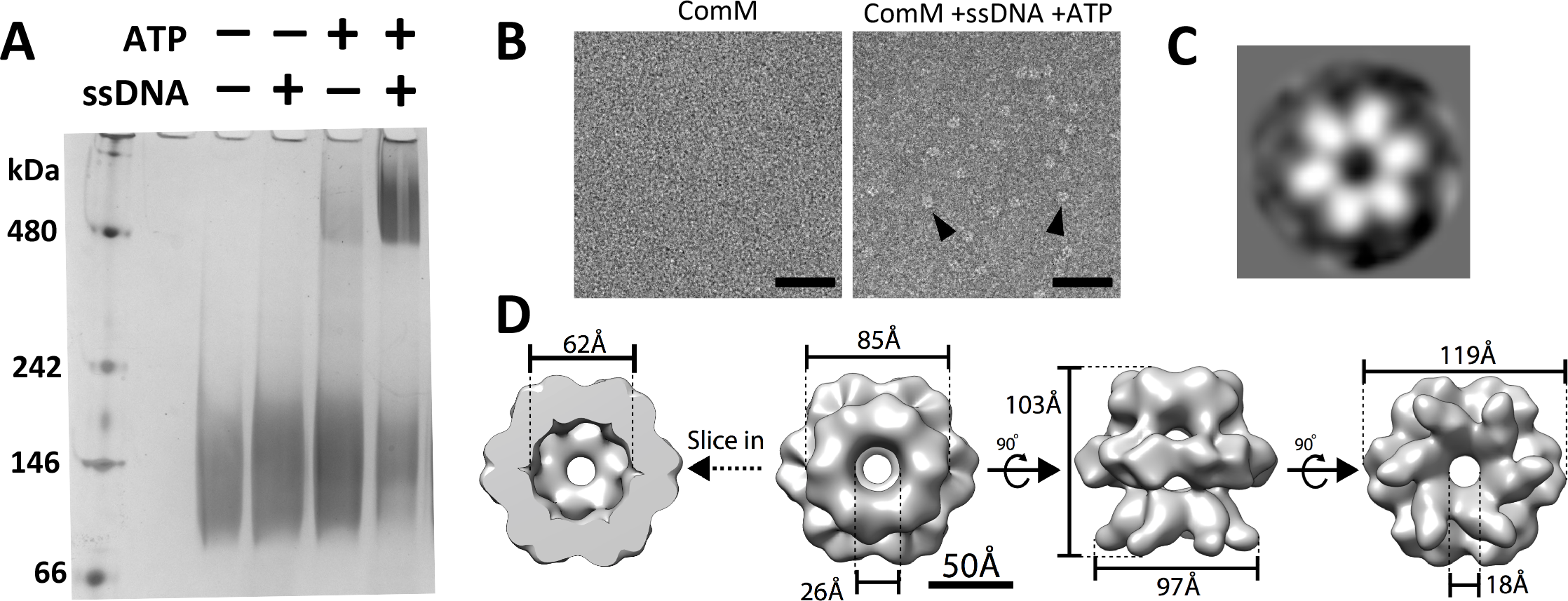
*ComM hexamerizes in the presence of ATP and ssDNA*. (**A**) Blue-native PAGE assay of purified ComM in the indicated conditions. (**B**) Negative stain EM of purified ComM under the indicated conditions. Representative ring-like densities observed in the presence of 5 mM ATP and 5 μM ssDNA are indicated by black arrows. Scale bar = 50 nm. (**C**) A representative 2-D class average of the ring-like densities observed by negative stain EM reveals a hexameric complex. (**D**) 3-D reconstruction (~13.8 Å resolution) of the ring complex imposing C6 symmetry.

### ComM binds ssDNA and dsDNA in the presence of ATP

Because ATP was required for oligomerization in the presence of ssDNA, we hypothesized that it would also be required for DNA binding. To test this, we mutated the conserved lysine in the Walker A motif of ComM, which in other AAA+ ATPases, promotes ATP binding (26). *In vivo*, we find that this mutation abrogates ComM function (**Fig. S5A**), while protein stability is maintained (**Fig. S2**). To determine if ATP binding was required for ComM to bind ssDNA, we performed electrophoretic mobility shift assays (EMSAs) using purified ComM and ComM^K224A^ *in vitro*. We find that WT ComM binds both ssDNA and dsDNA in an ATP-dependent manner (**Fig. S5B-D**). Consistent with this, ComM^K224A^ displays greatly reduced ssDNA binding even in the presence of ATP (**Fig. S5B**). Also, using non-labeled competitor DNA of differing lengths in EMSAs, we found that ComM preferentially bound ssDNA >60bp in length. Taken together, these results indicate that ATP is important for ComM to bind DNA and function during natural transformation.

### ComM has helicase activity in vitro

Thus far, our data suggest that ComM may play a role in branch migration *in vivo*. Some branch migration factors (e.g., RuvAB and RecG (32,33)) display helicase activity. So next, we tested the helicase activity of purified ComM *in vitro*. We observed enzymatic unwinding of a forked DNA substrate with increasing concentrations of ComM. Assuming that ComM hexamers are the active oligomeric form, the helicase activity had an apparent K_M_ of 50.8 nM (**Fig. 4A**). As expected, the purified ATP binding mutant ComM^K224A^ did not display helicase activity in this assay compared to WT ComM and Pif1, a previously characterized helicase (34) that served as a positive control in these assays (**Fig. 4B**). Furthermore, the non-hydrolysable ATP analog ATPγS inhibited helicase activity, which is consistent with ATP hydrolysis being required for ComM function (**Fig. 4B**).

**Fig. 4.**
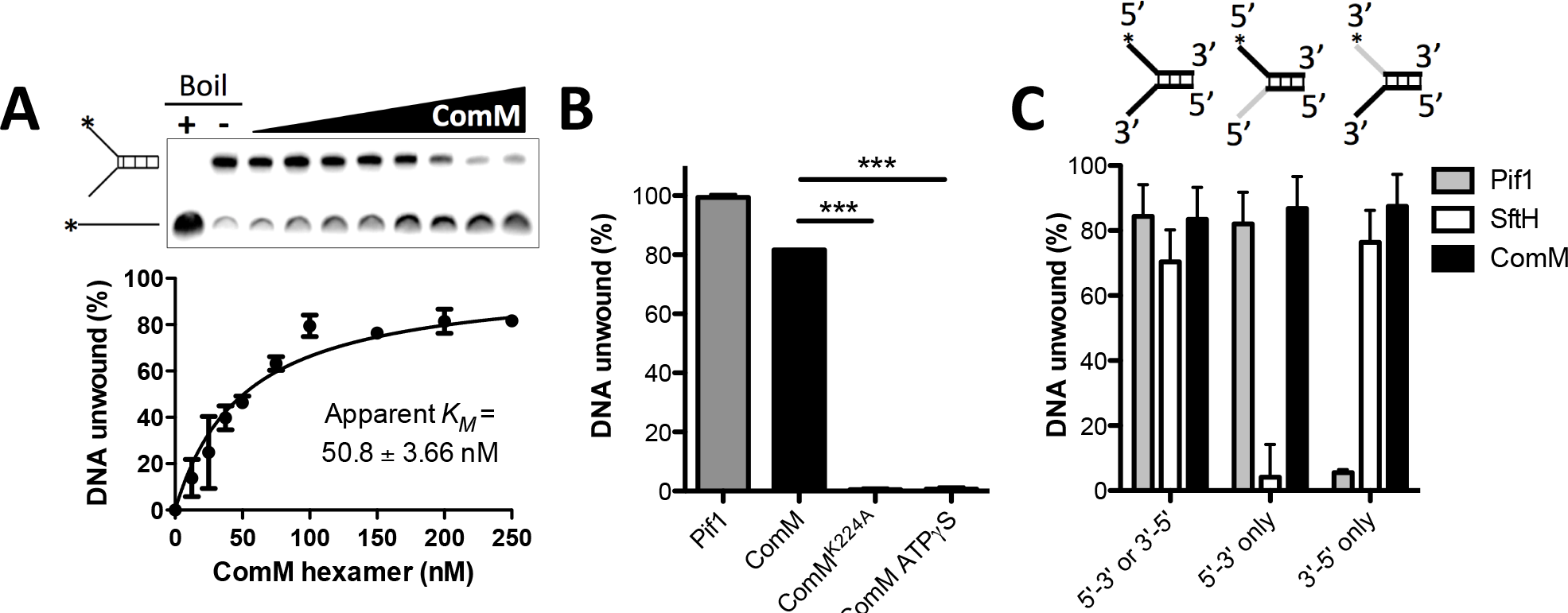
*ComM exhibits helicase activity* in vitro. (**A**) A representative forked substrate helicase assay with increasing concentrations of purified ComM. This forked substrate has 20 bp of annealed sequence and 25 bp tails. Concentrations of ComM used (in hexamer) were 0, 10, 25, 37.5, 50, 75, 100, 150, 200, and 250 nM. Images were quantified and plotted as indicated. (**B**) Helicase assay using forked DNA substrate with the indicated purified protein (100 nM Pif1 and 250 nM ComM / ComM^K224A^ hexamer) in the presence of 5 mM ATP (Columns 1, 2, and 3) or ATPγS (Column 4). (**C**) Helicase assays using forked DNA substrates that accommodate enzymes of either directionality or that can only be unwound by 5’ to 3’ or 3’ to 5’ activity. Directional substrates contained one ssDNA tail that is inverted relative to the remainder of the strand (inverted portion indicated in gray on the schematic above bars), thus, preventing helicase activity in one direction. Substrates were incubated with 100 nM purified ComM (hexamer), 10 nM Pif1, or 50 nM SftH. All data are shown as the mean ± SD and are the result of at least three independent replicates. *** = *p*<0.001.

Most helicases exhibit a preferred directionality (either 5’ to 3’ or 3’ to 5’). A forked DNA substrate, however, does not distinguish between these activities. To determine whether ComM had a preferred directionality, we tested helicase activity on forked substrates where one of the two tails is inverted (by a 3’-3’ linkage) relative to the remainder of the oligo (35). Thus, directional substrates either have two 5’ ends (to assess 5’ to 3’ directionality) or two 3’ ends (to assess 3’ to 5’ directionality). As controls in these assays, we used the unidirectional helicases Pif1 (a 5’ to 3’ helicase (34)) and SftH (a 3’ to 5’ helicase (36)). While Pif1 and SftH exhibited unidirectional helicase activity as expected, ComM unwound all of the substrates tested (**Fig. 4C**). Taken together, these data suggest that ComM is an ATP-dependent bidirectional helicase. Bidirectional activity of a single motor protein like ComM, while uncommon, is not unprecedented (37-39).

### ComM exhibits branch migration in vitro

While our data above clearly indicate that ComM is a hexameric helicase, not all helicases can promote branch migration. To more formally test whether ComM can promote branch migration, we performed *in vitro* branch migration assays using short 3-stranded substrates that more closely resemble the junctions of the D-loop that form during natural transformation (**Fig. S6A** and **S6B**)(40). These substrates contained a small region of heterology (indicated by the grey box), which prevents spontaneous branch migration as previously described (40). Using these substrates, we observed resolution of both the 5’ to 3’ and 3’ to 5’ substrates (**Fig. S6C** and **D**), which is consistent with the bidirectional helicase activity of ComM. This activity was inhibited when ComM was incubated with AMP-PNP, a nonhydrolyzable analog of ATP (Fig. **S6C** and **D**). And no activity was observed with ComM^K224A^ even if incubated with ATP (Fig. **S6C** and **D**). Thus, the branch migration activity observed is ATPase-dependent. It is formally possible that resolution of these short 3-stranded products was the result of only helicase activity (via the sequential removal of the unlabeled strand followed by the labeled strand or by unwinding of just the labeled strand). To address this, we also performed helicase assays using forked substrates that are derived from the oligos used to make the three-stranded branch migration substrates. This analysis indicated that ComM helicase activity on the forked substrates was markedly less efficient than its activity on the 3-stranded substrates (compare **Fig. S6C-D** to **S6E-F**), which suggests that the activity observed on the 3-stranded substrates is the result of *bona fide* branch migration and not simply helicase activity. The forked substrates used in this assay have 60 bp of annealed sequence (**Fig. S6E-F**), while the substrates previously used to test helicase activity only had 20 bp of annealed sequence (**Fig. 4A**). Thus, reduced activity on the forked DNA substrates in this assay is likely attributed to poor processivity for ComM helicase activity.

While the short 3-stranded substrates used above suggest that ComM possesses branch migration activity, we further tested 3-stranded branch migration using long DNA substrates. Long 3-stranded substrates were generated by RecA-mediated strand exchange between circular single-stranded (CSS) and linear double-stranded (LDS) φX174 DNA (23,41). Strand exchange reactions were allowed to proceed until the majority of the LDS was reacted with the CSS to form intermediates (INT = joint molecules that have not completed strand exchange) or nicked product (NP = the product formed upon complete strand exchange). Reactions were then deproteinated, and the resulting DNA was used to assess resolution of the long 3-stranded INT via branch migration (23,41). We found that ComM could efficiently drive the reaction in both directions, resolving the INT structures into both NP and LDS (**Fig. 5**), while reactions where ComM^K224A^ was added showed no activity. These results are consistent with ComM promoting branch migration of these long DNA substrates. Together, these results indicate that ComM exhibits branch migration on 3-stranded junctions *in vitro* on substrates that are likely similar to the structures formed during natural transformation *in vivo*.

**Fig 5.**
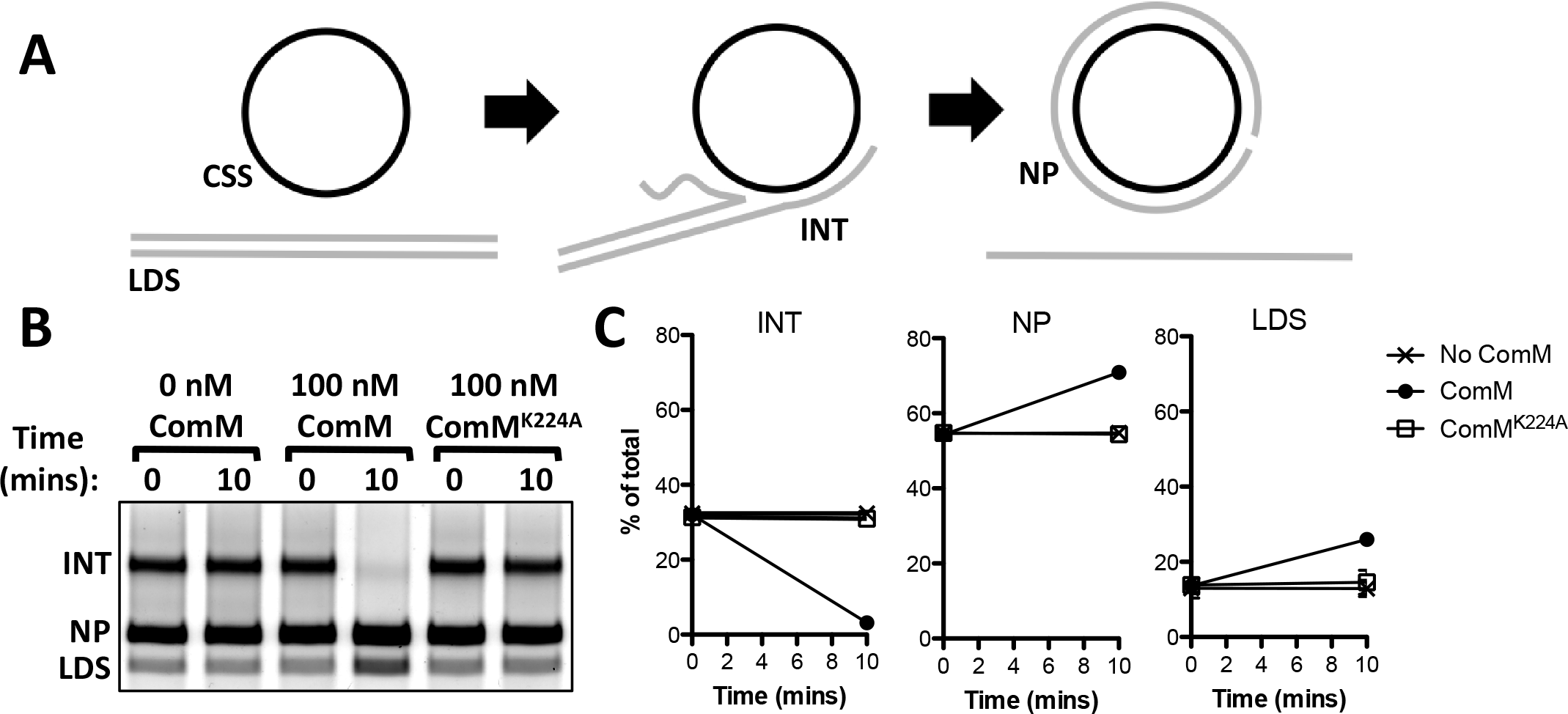
*ComM exhibits 3-stranded branch migration activity on long DNA substrates* in vitro. (**A**) Schematic for RecA-mediated strand exchange between linear double stranded PhiX174 (LDS) and circular single-stranded PhiX174 (CSS), which results in the formation of intermediates (INT) that can be resolved to nicked product (NP) if strand exchange commences to completion. Strand exchange reactions were deproteinated prior to complete strand exchange, and the resulting DNA was used to assess branch migration-dependent resolution of intermediate structures (INT). (**B**) Representative gel where deproteinated intermediates were incubated with the proteins indicated. (**C**) Three independent replicates of the assay described in **B** were quantified, and the relative abundance of the INT, NP, and LDS are shown as the mean ± SD.

### ComM is broadly conserved

Next, we assessed how broadly conserved ComM was among bacterial species. Homologs of ComM are found almost ubiquitously among Gram-negative species, including all known Gram-negative naturally competent microbes (**Fig. 6** and **Fig. S7**). Among this group, only select species lacked a ComM homolog, suggesting that loss of ComM occurred relatively recently (**Fig. 6** and **Fig. S7**). By contrast, we see a pervasive lack of ComM homologs among species within the Bacilli and Mollicutes, (**Fig. 6** and **Fig. S7**), suggesting that ComM may have been lost in a common ancestor for these two Classes. Interestingly, all known naturally competent Gram-positive species fall within the Bacilli group, indicating that branch migration during natural transformation must occur via a ComM-independent mechanism in these microbes.

**Fig. 6.**
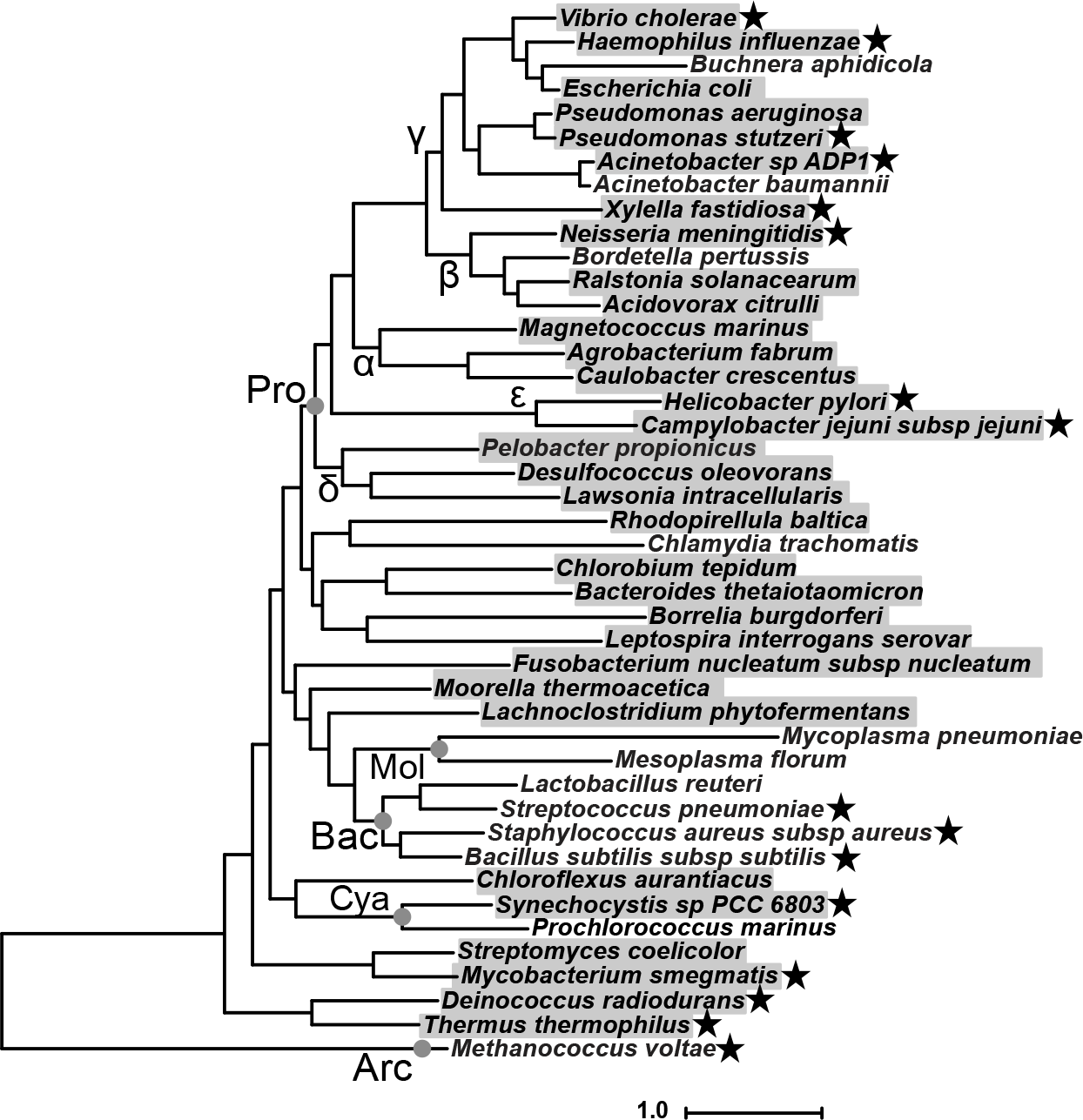
*ComM is broadly conserved*. Estimated maximum likelihood phylogeny of diverse species based on a concatenated alignment of 36 conserved proteins identified from whole genome sequences. Species with an identified ComM homolog are highlighted in gray. Competent species are designated by a star next to the species name. Major taxa are labeled along their nodes. Pro: Proteobacteria (Greek letters indicate subdivisions); Bac: Bacilli; Mol: Mollicutes; Cya: Cyanobacteria; Arc: Archaea. Scale bar indicates distance.

### ComM promotes branch migration in Acinetobacter baylyi

Because ComM is broadly conserved among Gram-negative naturally competent microbes, next, we tested its role during natural transformation in the model competent species *A. baylyi* ADP1. As in *V. cholerae*, a *comM* mutant of *A. baylyi* displayed greatly reduced rates of natural transformation when using linear tDNA (**Fig. 7A, left**). Also, this mutant displayed reduced comigration of linked genetic markers, consistent with *comM* playing a role in branch migration (**Fig. 7A, right**). Additionally, we observed that ComM is required for integration of tDNA containing larger regions of heterologous sequence (**Fig. 7B**). These data are consistent with what was observed in *V. cholerae* (**Fig. 2**) and suggests that ComM is a conserved branch migration factor important for natural transformation in diverse Gram-negative bacterial species.

**Fig. 7.**
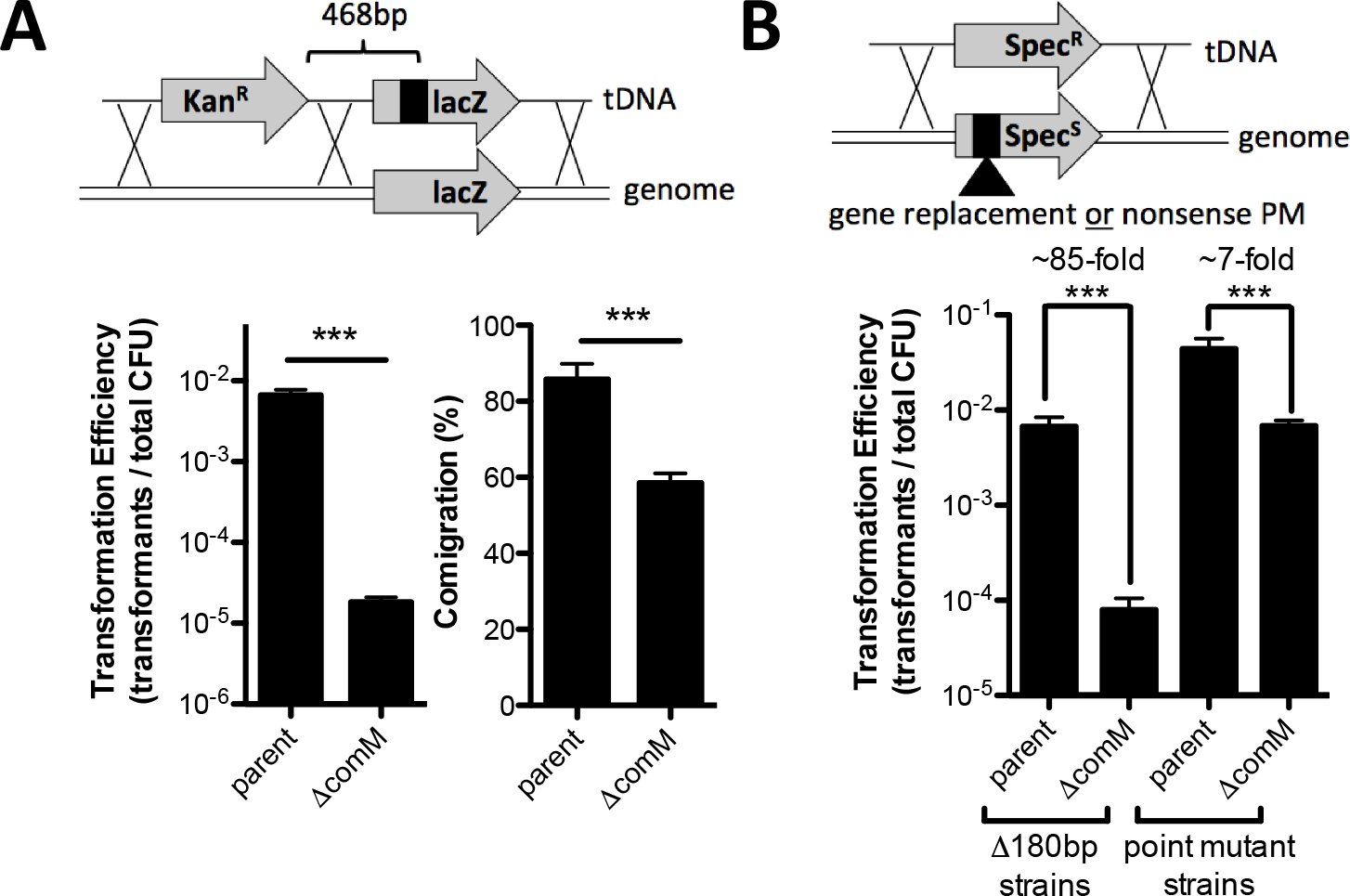
*ComM promotes branch migration in* Acinetobacter baylyi. (**A**) Transformation assay of *A. baylyi* using a linear tDNA product with linked genetic markers. (**B**) Transformation assay performed in Spec^S^ strains with tDNA that would revert the strain to Spec^R^. Integration of the marker would either repair a point mutation or delete 180 bp of genomic sequence. All strains in **A** and **B** contain *mutS* mutations to prevent MMR activity. In schematics above bar graphs, X’s denote possible crossover points for homologous recombination. All data are shown as the mean ± SD and are the result of at least three independent biological replicates. *** = *p*<0.001.

### RadA is not required for natural transformation in V. cholerae

While ComM is broadly conserved, it is absent in all of the known Gram-positive naturally competent species (**Fig. 6**). In the Gram-positive *Streptococcus pneumoniae*, mutants of *radA* (also known as *sms*) display reduced rates of natural transformation but are not affected at the level of tDNA uptake, similar to what is observed for *V. cholerae comM* mutants in this study (42) and in *H. influenzae* (9). Similar results are also seen in *radA* mutants in *Bacillus subtilis* (43). *E. coli* RadA has recently been shown to promote branch migration during RecA-mediated strand exchange (23). Also, it was very recently demonstrated that RadA is a hexameric helicase that promotes bidirectional D-loop extension during natural transformation in *S. pneumoniae* (38). Interestingly, *radA* is broadly conserved and both *V. cholerae* and *A. baylyi* contain *radA* homologs (VC2343 and ACIAD2664, respectively). RadA, however, is not required for natural transformation in *V. cholerae* (**Fig. S8**). Also, preliminary Tn-seq data from our lab indicates that RadA is not important for natural transformation in *A. baylyi*. Thus, it is tempting to speculate that in Gram-positive competent species, RadA carries out the same function that ComM plays during natural transformation in Gram-negative species.

### ComM is not required for DNA repair

ComM homologs are also found among diverse non-competent species (**Fig. 6**). Moreover, many bacterial helicases are implicated in promoting branch migration during other types of homologous recombination, including during DNA repair. So next, we wanted to determine if ComM also plays a role in DNA repair independent of its role during natural transformation. ComM is poorly expressed in the absence of competence induction in *V. cholerae* (**Fig. S2**). Thus, to test the role of ComM in DNA damage, we tested survival of P_*tac*_-*tfoX* and P_*tac*_-*tfoX* Δ*comM* strains under competence inducing conditions (i.e. ectopic expression of TfoX). A WT strain and a *recA* mutant were also included in these assays as controls. DNA damage was tested using methyl methanesulfonate (MMS - methylates DNA / stalls replication forks) and mitomycin C (MMC – alkylates DNA / generates interstrand DNA crosslinks)(44,45). As expected, the *recA* mutant was more sensitive to these treatments compared to the WT, consistent with a critical role for homologous recombination in DNA repair (**Fig. S9**) (46,47). The P_*tac*_-*tfoX* Δ*comM* mutant, however, was as resistant to these DNA damaging agents as the P_*tac*_-*tfoX* strain, indicating that this branch migration factor either does not play a role during DNA repair or that the activity of this protein is redundant with other branch migration factors in the context of repair (**Fig. S9**). Furthermore, induction of competence (via ectopic expression of TfoX) showed little to no difference in DNA damage repair compared to the WT, indicating that natural competence plays a limited role in DNA repair in *V. cholerae* under the condition tested (**Fig. S9**).

## DISCUSSION

Natural transformation is an important mechanism of horizontal gene transfer in bacterial species. It is dependent upon activation of bacterial competence or the ability to bind and take up exogenous DNA. Altogether, our *in vivo* and *in vitro* data elucidate a role for ComM as a helicase / branch migration factor that promotes the integration of tDNA during natural transformation. In our model, integration is initiated by RecA-mediated strand invasion and formation of a D-loop that generates a three-stranded intermediate structure (**Fig. 8**). Through its bidirectional helicase and/or branch migration activity, ComM then likely promotes expansion of the D-loop, which enhances the integration of tDNA into the genome (**Fig. 8**). Following branch migration, the junctions are resolved to mediate stable integration of tDNA by an unresolved mechanism.

**Fig. 8.**
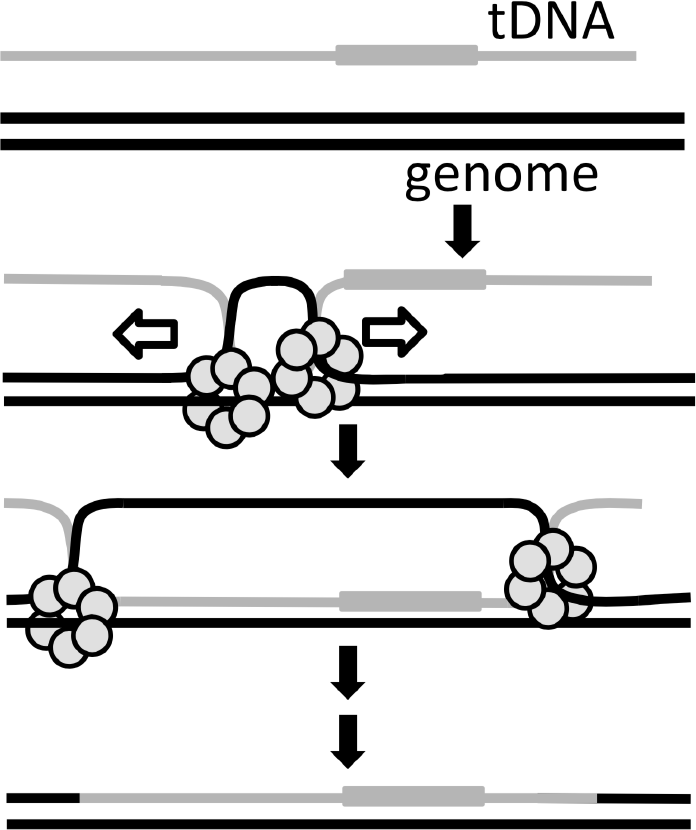
*Proposed model for the role of ComM during natural transformation*. ComM is shown as a hexameric ring that promotes integration of tDNA via its bidirectional helicase and/or branch migration activity. This can support integration of tDNA with a heterologous region, which is indicated by a gray box.

Our data suggest that ComM is important for the incorporation of heterologous sequences. The main drivers for evolution and maintenance of natural transformation in bacterial species are heavily debated. One model suggests that this process is largely for enhancing genetic diversity, while another hypothesis is that natural transformation evolved as a mechanism for acquisition of DNA as a nutrient (48,49). These processes are not mutually exclusive. Because ComM affects only the integration of heterologous tDNA (and not its uptake), however, the activity of this protein supports a role for natural transformation in adaptation and evolution through the acquisition of novel genetic material. Other competence genes that are involved specifically in homologous recombination (e.g., *dprA*) also support this hypothesis.

Our *in vitro* data suggest that ComM forms a hexameric ring structure in its active state similar to that of eukaryotic MCM2-7 and bacterial DnaB (31,50). Our 3D reconstruction reveals a three-tiered barrel-like complex with a ~26-Å pore and ~18-Å pore on the top and bottom, respectively. We propose that ComM either forms around, or is loaded onto the displaced single strand of the native genomic DNA and acts as a wedge to dissociate these strands. Alternatively, dsDNA may enter the 26-Å pore, be unwound in the central channel of the complex with single strands bring extruded through the side channels that are evident between the tiers of the barrel structure. Such side channels are not uncommon among hexameric AAA+helicases (*e.g*., SV40 T-antigen (51,52) and the archaeal MCM (28,53)) and have been hypothesized to be exit channels for extruded ssDNA. The pore sizes observed in our reconstruction appear to be consistent with those found in other ring helicases, however, it has been shown that the size of the opening and channel can change depending on the nucleotide bound (ATP vs. ADP) (51,54). Future work will focus on characterizing the ComM structure bound to ADP and AMP-PNP, which may help inform the structural changes associated with the catalytic cycle of this hexameric helicase.

Our phylogenetic analysis indicates that ComM is broadly conserved in bacterial species, and is largely, only excluded from the Bacilli and Mollicutes. There is also, however, evidence for isolated examples for ComM loss in species that fall outside of these two groups, which suggests that these have occurred relatively recently. Some of these represent obligate intracellular pathogens or endosymbiots, which commonly contain highly reduced genomes (e.g. *Buchnera* and *Chlaymdia*) (55,56). Another example of recent ComM loss is among *Prochlorococcus* species, which are related to other naturally competent cyanobacteria (e.g. *Synechococcus* and *Synechocystis spp*.). Interestingly, we found that *Prochlorococcus* lacked many of the genes required for natural transformation (e.g. *dprA*, *comEA*, *comEC*, etc.) (57), indicating that competence may have been lost in this lineage of cyanobacteria. Another notable example of ComM loss is in *Acinetobacter baumanii*, which is an opportunistic pathogen that is closely related to *A. baylyi*. Many strains of *A. baumanii*, including the one analyzed here (ATCC 17978), contain an AbaR-type genomic island integrated into *comM* (58,59). Strains that contain this horizontally transferred genomic island display low rates of natural transformation while those that lack it display higher rates of natural transformation (60), indicating that AbaR island-dependent inactivation of ComM may inhibit this mechanism of horizontal gene transfer in *A. baumanii*. This represents another example in a growing list of horizontally acquired genomic elements that inhibit natural transformation (61-63). Our phylogenetic analysis also indicates that ComM is highly conserved among non-competent bacterial species. This suggests that ComM may have a function outside of natural transformation. Alternatively, it is possible that many of these species are capable of natural transformation, however, the conditions required for competence induction have not yet been identified. Indeed, the inducing cue for natural transformation in *V. cholerae* was only discovered in 2006 (2).

Our data suggest that branch migration in Gram-positive and Gram-negative species has diverged in their dependence on distinct factors (RadA and ComM, respectively). The tract length of DNA recombined into *S. pneumoniae* during natural transformation is ~2.5 kb on average (64). In *H. influenza*e, the mean recombination tract length is ~14 kb (65). Thus, compared to RadA-dependent branch migration in Gram-positive species, ComM may facilitate the integration of more tDNA in Gram-negative species. Other proteins that impact DNA integration during natural transformation, however, may confound this overly simplified comparison.

ComM is not essential for natural transformation because transformants are still observed in a *comM* mutant. This suggests that other proteins may be involved in promoting the integration of tDNA in the absence of this branch migration factor. Also, our data suggest that these alternative branch migration factors are less efficient at incorporating tDNA with sequence heterology compared to ComM in both *V. cholerae* and *A. baylyi*. This role could be carried out by another helicase. Candidates include RuvAB, RecG, and PriA, which have all previously been implicated in branch migration (32,33,66). In fact, the primasome helicase PriA is essential for natural transformation in *Neisseria gonorrhoeae*. This may be due to a role in branch migration, however, PriA is also essential for restarting stalled replication forks and *priA* mutants have a severe growth defect (67). Also, it has been proposed that PriA helicase activity may facilitate the uptake of tDNA through the inner membrane (68). A *recG* homologue in *Streptococcus pneumoniae* (*mmsA*) was shown to have a mild effect on transformation when deleted (69). RecG reverses stalled replication forks and promotes branch migration opposite to the direction of RecA-mediated strand exchange in *E. coli* (70,71). Also, RecG acts on three-strand intermediates *in vitro*, which could give credence to involvement in natural transformation where such a structure is formed (41). Because RecG works counter to the direction of RecA-mediated strand exchange, however, it is currently unclear how RecG may play a role in natural transformation in the presence of ComM. Another possibility is that RecA, which has inherent ATP-dependent unidirectional branch migration activity (72,73), could promote branch migration independent of other canonical branch migration factors. Future work will focus on identifying genetically interacting partners of ComM to uncover the role of additional factors required for efficient integration of tDNA during natural transformation.

In addition to a AAA+ ATPase domain, ComM also contains two magnesium (Mg) chelatase domains. Mg chelatase domain-containing proteins have only previously been implicated in inserting Mg into protoporphyrin rings in photosynthetic organisms (74). While it is currently unclear how these domains participate in ComM function, to our knowledge, this study is the first to indicate a function for a Mg-chelatase domain-containing protein in a non-photosynthetic organism.

In conclusion, the results from this study strongly support that ComM enhances natural transformation by promoting ATP-dependent bidirectional helicase activity and/or branch migration activity, which allows for the efficient integration of heterologous tDNA. Furthermore, our data in *V. cholerae* and *A. baylyi* as well as previous work from *H. influenzae* indicate that this branch migration factor is a broadly conserved mechanism for integration of tDNA in diverse naturally transformable Gram-negative species.

## ACKNOWLEDGEMENTS

This work was supported by US National Institutes of Health Grant AI118863 to A.B.D., an Indiana University Collaborative Research Grant to M.L.B., an American Cancer Society Research Scholar grant (RSG-16-180-01-DMC) to M.L.B., startup funds from the Indiana University College of Arts and Sciences to ABD, and DTK was supported by NIH grant R35 GM122556 to Yves Brun.

## Supplementary figures

**Fig. S1.**
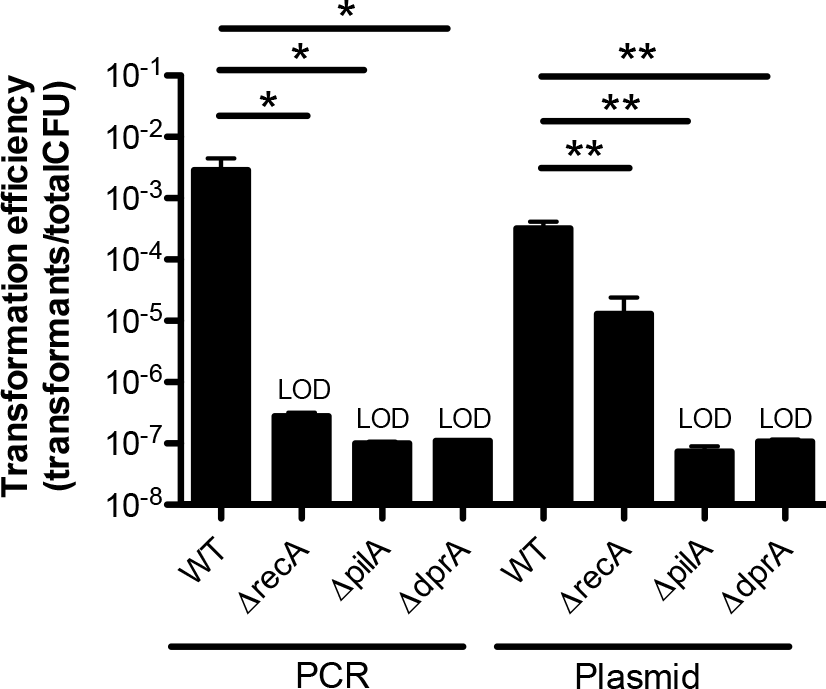
*Uptake of plasmid DNA is independent of recombination*. Chitin-dependent transformation assay with the indicated strains using either linear PCR product or plasmid as tDNA. All data are shown as the mean ± SD and are the result of at least three independent biological replicates. * = *p*<0.05, ** = *p*<0.01, and LOD = limit of detection.

**Fig. S2.**
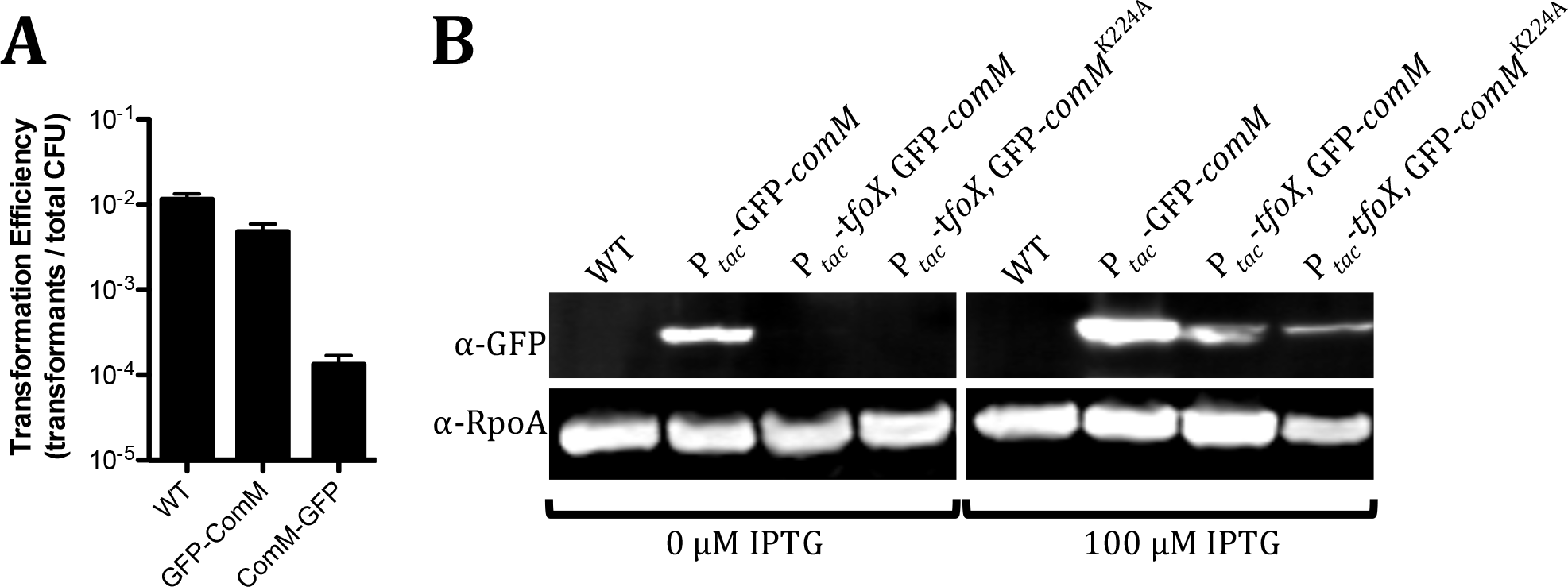
*N-terminal ComM fusions are functional*. (**A**) Chitin-dependent transformation assay with the indicated strains using linear PCR product as the tDNA. (**B**) Representative western blot to detect GFP-ComM and RpoA (loading control) in the indicated strains grown in the presence or absence of 100 μM IPTG. Blot indicates that GFP-*comM* and GFP-*comM*K224A at the native locus are induced when TfoX is ectopically expressed to induce competence. Also, this blot indicates that the P*tac*-GFP-*comM* construct is leaky and expressed in the absence of inducer. Data in **A** are shown as the mean ± SD and are the result of at least three independent biological replicates.

**Fig. S3.**
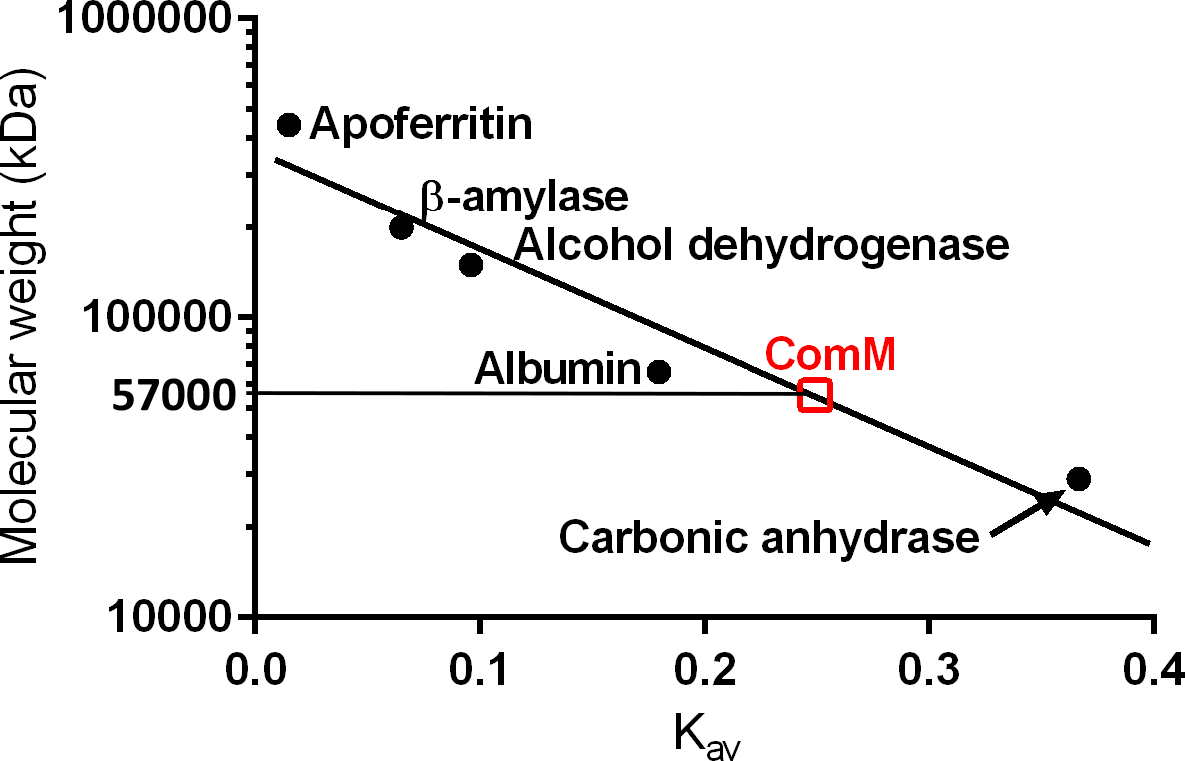
*ComM is monomeric in soluble form*. Purified StrepII-ComM was analyzed by gel filtration and compared to a set of protein standards to determine size.

**Fig. S4.**
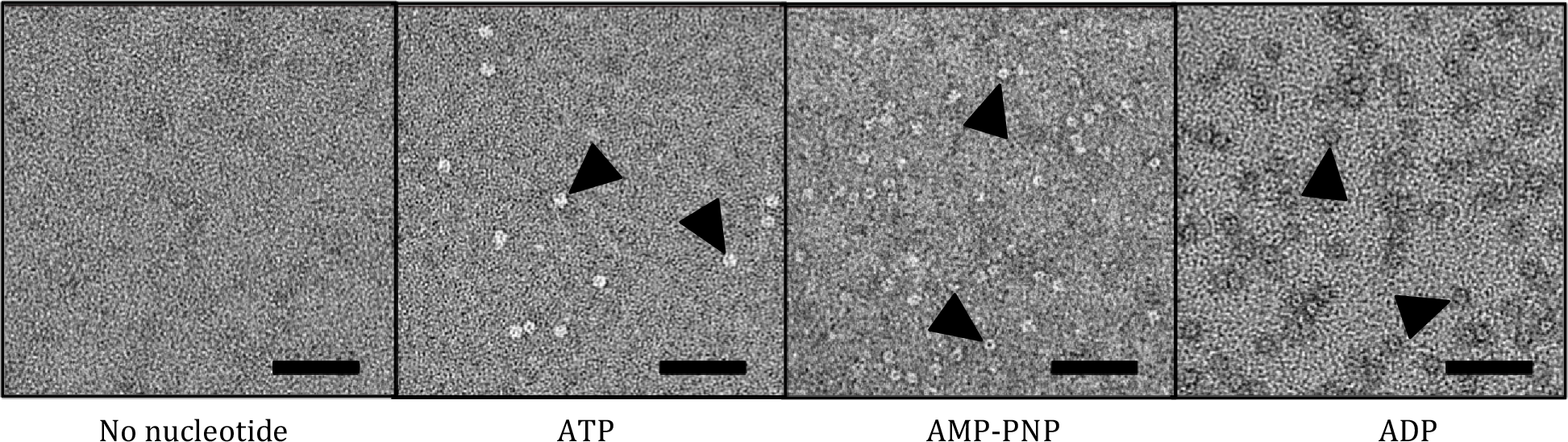
*ComM oligomerizes in the presence of ADP and AMP-PNP*. Negative stain EM of purified ComM incubated with 5 μM ssDNA and 5mM of the indicated nucleotide or nucleotide analog. Representative ring-like densities in these samples are indicated by black arrows. Scale bar = 50 nm.

**Fig. S5.**
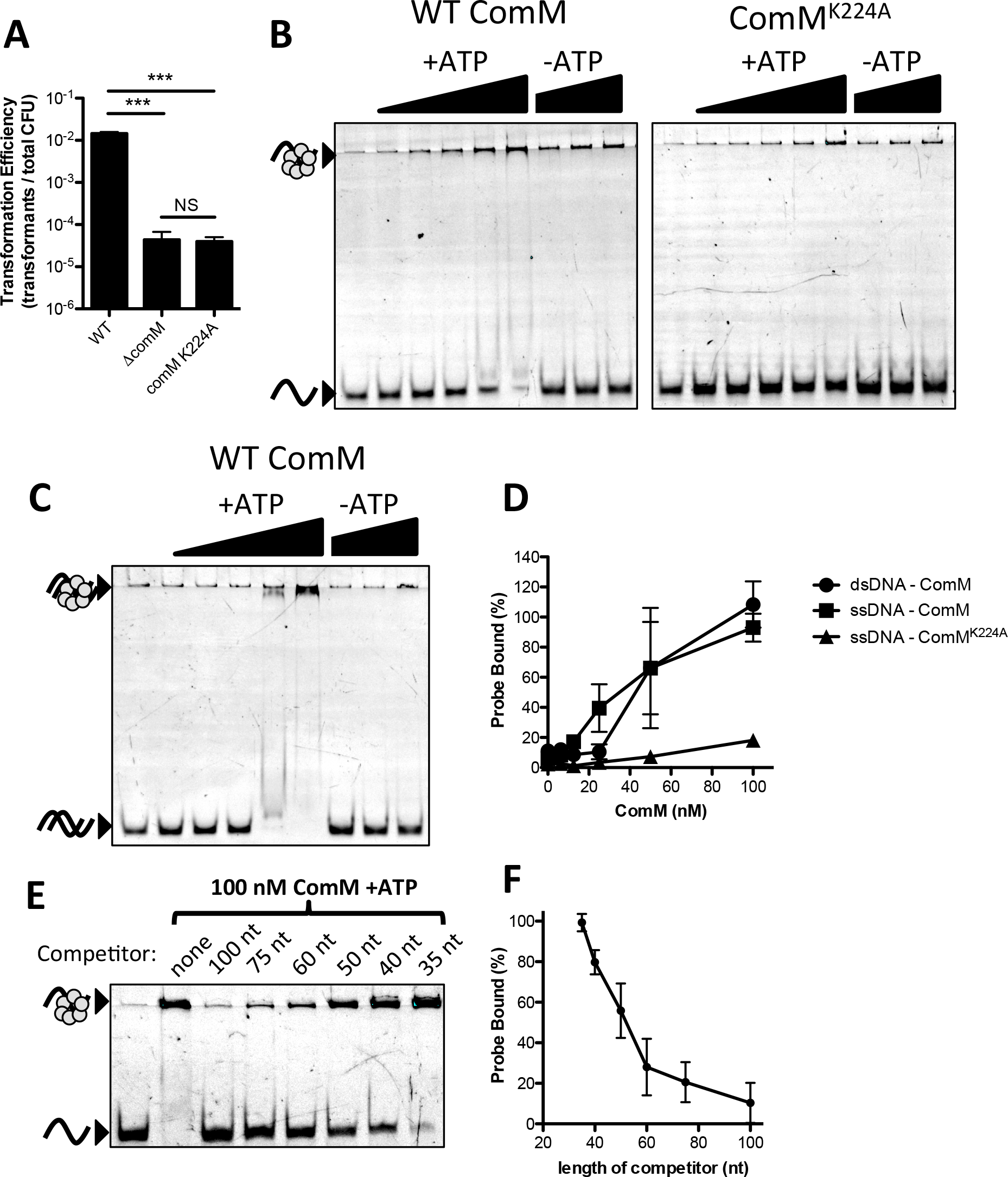
*ComM binds ssDNA in the presence of ATP*. (**A**) Chitin-dependent transformation assay of the indicated strains using a linear PCR product as tDNA. Data are shown as the mean ± SD and are the result of at least three independent biological replicates. *** = *p*<0.001, NS = not significant. (**B**) EMSA with purified ComM and ComMK224A and a ssDNA probe. Protein concentrations (of the hexamer) used in the presence of ATP (+ATP) were 0, 6.25, 12.5, 25, 50, 100 nM, and in the absence of ATP (-ATP) were 25, 50, and 100 nM. Bound probe is retained in the well due to the large size of the DNA-bound oligomeric complex. (**C**) Representative EMSA with purified ComM and dsDNA probe. All reactions were performed in the presence of ATP and the protein concentrations (of hexamer) used were the same as in **B.** (**D**) Replicate EMSAs from **B** and **C** were quantified and plotted as indicated. (**E**) A representative EMSA where binding was competed with cold competitor DNA of the indicated length. The labeled probe was a 100 nt poly-dT oligo. The cold competitor was of the length indicated (also poly-dT) and was added at 100X molar excess to the labeled probe. (**F**) Replicate EMSAs from **E** were quantified and plotted as indicated. Data in **B**, **C**, and **E** are representative of at least three independent experiments.

**Fig. S6.**
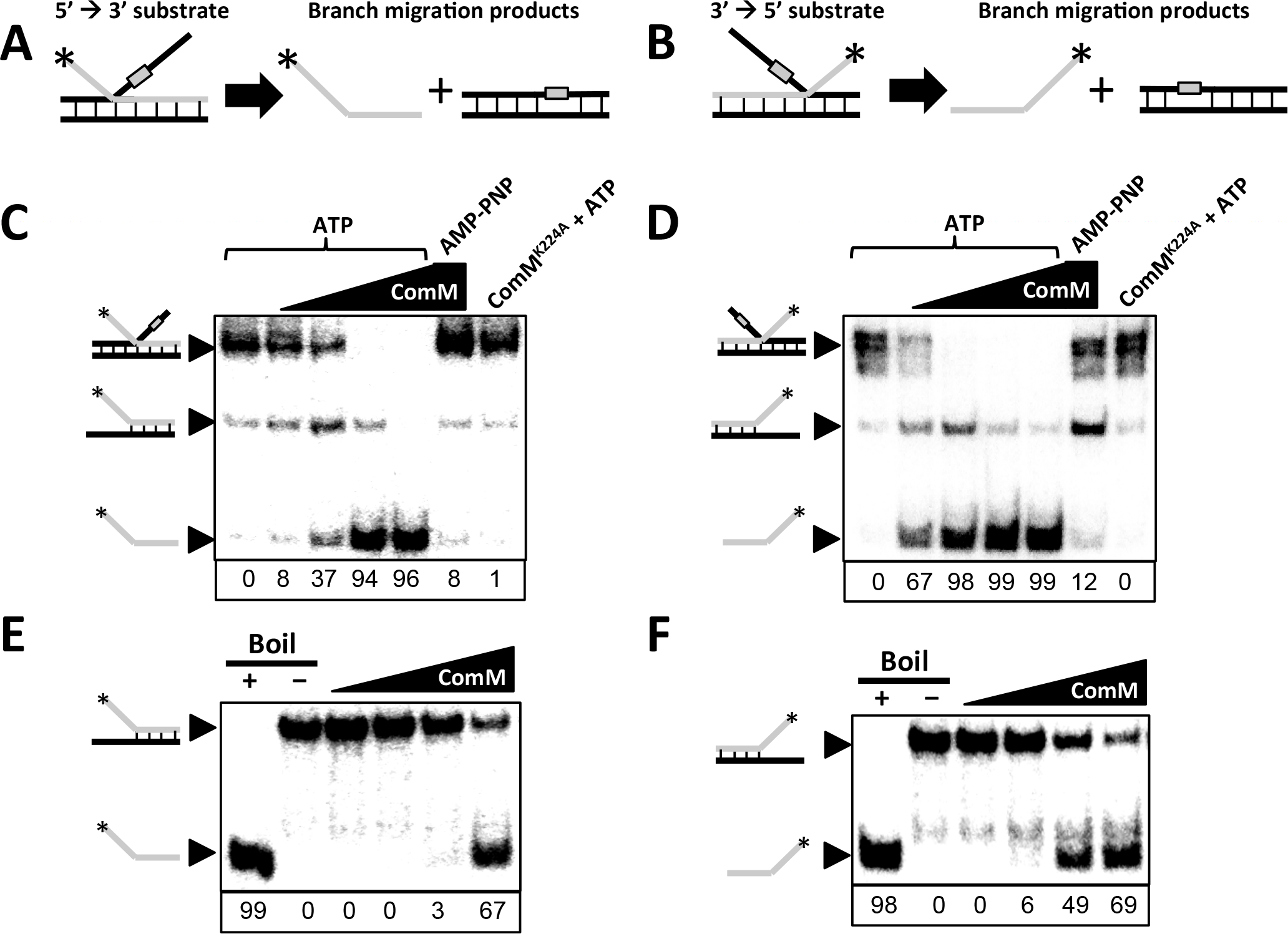
*ComM exhibits branch migration activity on short substrates* in vitro. Schematics of the substrates used to test (**A**) 5’→3’ and (**B**) 3’→5’ branch migration activity. The gray box indicates a region on the substrate that is not homologous to the complementary strand. This was introduced to prevent spontaneous branch migration. The labeled strand has 60 bp of annealed sequence and a 30 nt tail (**C** and **D**) Representative branch migration assays using the substrates described in **A** and **B**, respectively. Substrates were incubated with 0, 25, 50, 100, or 200 nM WT ComM hexamer or 200 nM ComMK224A as indicated. Reactions were incubated with ATP or AMP-PNP as indicated. The % of final branch migration product generated at each concentration of ComM is indicated below each lane. (**E** and **F**) Representative helicase assays using forked substrates derived from the same oligos used to generate the three-stranded branch described in **A** and **B**, respectively. Substrates were incubated with 0, 25, 50, 100, or 200 nM WT ComM hexamer in the presence of ATP. The % of unwound product generated at each concentration of ComM is indicated below each lane. All data are representative of at least three independent experiments.

**Fig. S7.**
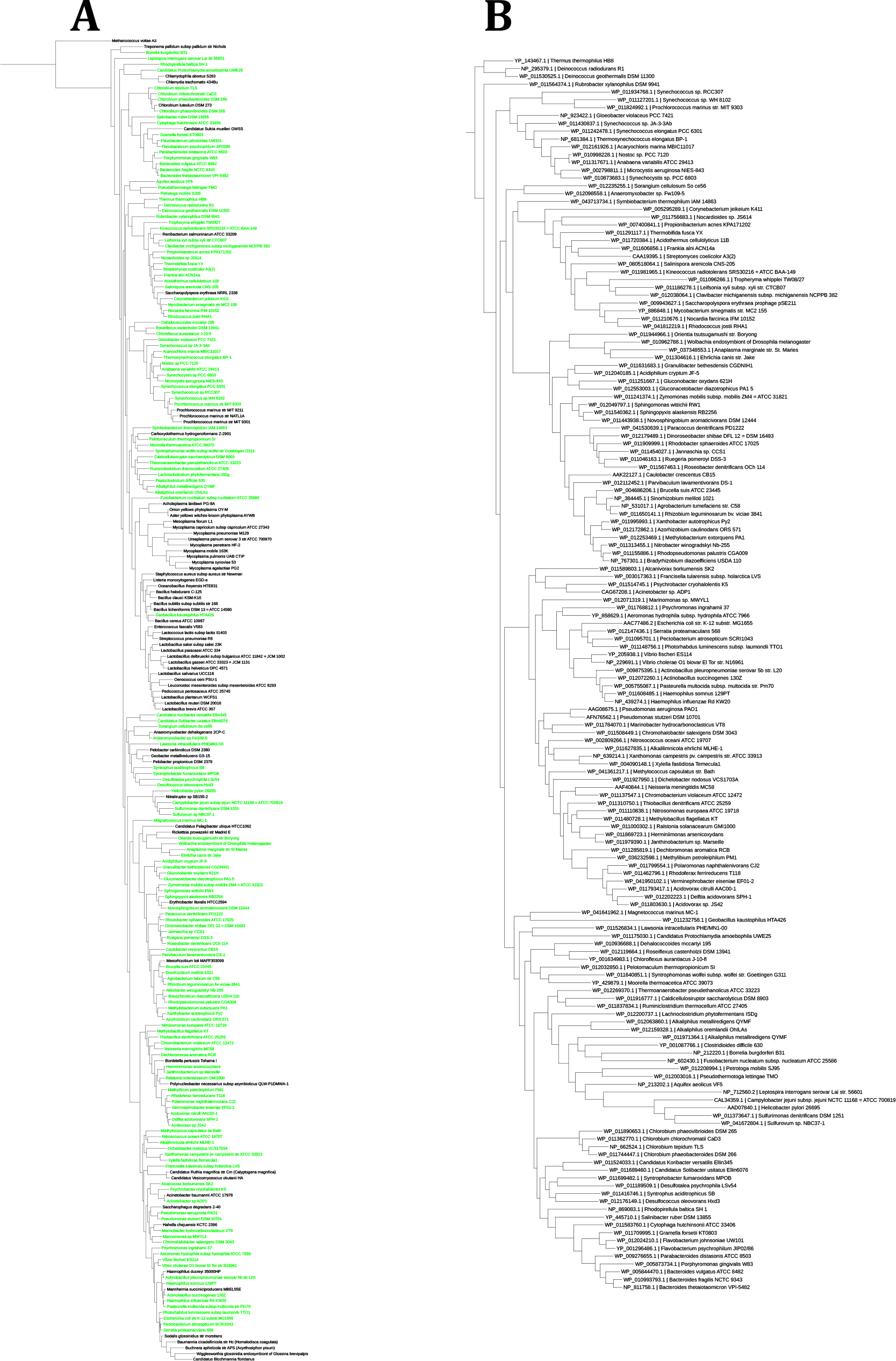
*ComM is broadly conserved*. (**A**) Phylogenetic trees of species based on a concatenated alignment of 36 conserved protein sequences. Green text indicates species with an identifiable ComM homolog. (**B**) Phylogenetic tree of ComM alleles.

**Fig. S8.**
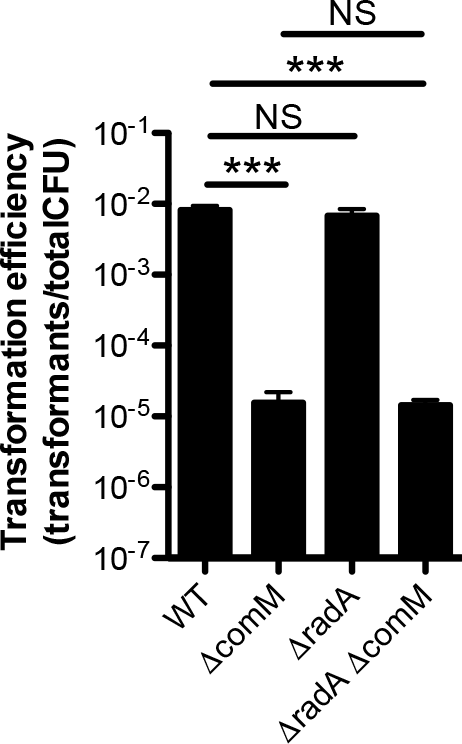
*RadA is not required for natural transformation in* V. cholerae. All strains contain P*tac*-*tfoX* mutations and were transformed via chitin-independent transformation assays using a linear PCR product as the tDNA. All data are shown as the mean ± SD and the result of 6 independent biological replicates. *** = *p*<0.001, NS = not significant.

**Fig. S9.**
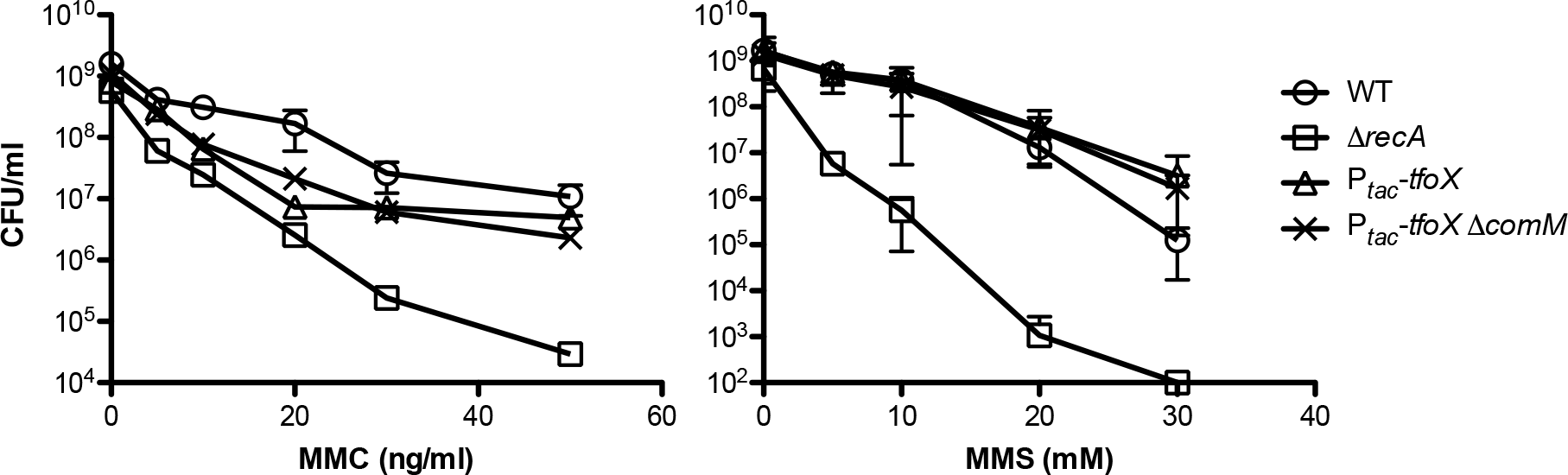
*ComM is not required for DNA repair*. Strains were treated with increasing doses of the DNA damaging agent indicated on the X-axis and then plated for viability. All data are shown as the mean ± SD and are the result of at least three independent biological replicates.

**Table S1.** – Strains used in this study.

| Strain Name in Manuscript | Genotype and Antibiotic Resistances | Description | Reference / strain# |
| --- | --- | --- | --- |
| WT | Sm<sup>R</sup> | Wildtype *V. cholerae* O1 El Tor strain used throughout this study | (1) / (SAD030) |
| **Strains Used for Transformation Assays** |  |  |  |
| $\Delta comM$ | $\Delta VC0032::Spec^R$ | Replacement of VC0032 with a spectinomycin resistance cassette | This Study (SAD083) |
| $P_{tac-tfoX}$ | VC1153 OE Kan<sup>R</sup> | OE Kan<sup>R</sup> = a fragment of the Tn10 transposon from pDL1093, including the *rrnB* antiterminator, *Ptac*, *LacI*, and Kan<sup>R</sup>. This fragment, and its use in overexpression of VC1153 (*tfoX*) is described in | (2) / (SAD061) |
| $\Delta comM P_{tac-tfoX}$ | VC0032::Spec<sup>R</sup>, VC1153 OE Kan<sup>R</sup> | Replacement of VC0032 with Spec<sup>R</sup> cassette in a VC1153 OE Kan<sup>R</sup> parent strain | This Study (SAD066) |
| $P_{tac-comM}$ | $P_{tac-comM}$ at *lacZ* locus Spec<sup>R</sup> | VC0032 fused with an IPTG inducible promoter at the *lacZ* locus | This Study (SAD1065) |
| $\Delta recA$ | $\Delta recA::Spec^R$ | Replacement of VC0543 with Spec<sup>R</sup> cassette | This Study (SAD081) |
| $\Delta pilA$ | $\Delta pilA::Spec^R$ | Replacement of VC2423 with Spec<sup>R</sup> cassette | This Study (SAD780) |
| $\Delta dprA$ | $\Delta dprA::Spec^R$ | Replacement of VC0048 with Spec<sup>R</sup> | This Study (SAD079) |
| $P_{tac-comM}, \Delta comM$ | $P_{tac-comM}$ at *lacZ* locus Spec<sup>R</sup>, $\Delta VC0032::Amp^R$ | VC0032 fused with an IPTG inducible promoter at the *lacZ* locus in a $\Delta comM$ parent strain | This Study (SAD1066) |
| $\Delta 29bp$ (Tm<sup>S</sup>) | $\Delta VC1807::TmR^*\Delta 29bp$, $\Delta mutS$ MuGENT edit, LPQEN KanR | $\Delta VC1807::TmR^*\Delta 29bp$, $\Delta mutS$ MuGENT edit, LPQEN KanR; NOT TmR resistant | This Study (TND0226/ SAD1321) |
| $\Delta 29bp$ (Tm<sup>S</sup>) $\Delta comM$ | $\Delta comM::CarbR$, ($\Delta VC1807::TmR^*\Delta 29bp$, $\Delta mutS$ MuGENT edit, LPQEN KanR) | VC0032 deletion in a Tm<sup>S</sup> parent strain | This Study (TND0229/ SAD1322) |
| Point mutant (Tm<sup>S</sup>) | $\Delta mutS$ MuGENT edit, LPQEN::SpecR, $\Delta VC1807::TmR^*TI$ | $\Delta mutS$ MuGENT edit, LPQEN::SpecR, $\Delta VC1807::TmR^*TI$; $\Delta VC1807::TmR^*TI$ is the TmR cassette at VC1807, except it has a transition point mutation that introduces a premature stop codon. | This Study (TND0220/ SAD1323) |
| Point mutant (Tm<sup>S</sup>), $\Delta comM$ | $\Delta mutS$ MUGENT edit, $\Delta comM::CarbR$, LPQEN::KanR, $\Delta VC1807::TmR^*TI$ | $\Delta mutS$ MUGENT edit, $\Delta comM::CarbR$, LPQEN::KanR, $\Delta VC1807::TmR^*TI$; $\Delta VC1807::TmR^*TI$ is the TmR cassette at VC1807, except it has a transition point mutation that introduces a premature stop codon. | This Study (TND0221/ SAD1324) |
| $comM^{K224A}$ | LPQEN::Kan<sup>R</sup>, $comM^{K224A}$ | K to A residue substitution disrupts ATP binding. FRT Kan cassette following the LPQEN amino acid sequence in *lacZ* (VC2338) used for selection during co-transformation. | This Study (SAD1026) |
| *gfp-comM* | *gfp-comM* | N-terminal GFP-comM at native locus | This Study (SAD924) |
| *comM-gfp* | *comM-gfp* | C-terminal comM-GFP at native locus | This Study (SAD925) |
| parent (ADP1) | ΔACIAD1551::P<sub>tac</sub>-*lacZ*, ΔmutS::Spec<sup>R</sup> | lacZ introduced into a defunct transposase (neutral gene = ACIAD1551), mutS deleted and replaced with Spec<sup>R</sup> (ACIAD1500) | This Study (TND0137/ SAD1325) |
| ΔcomM (ADP1) | ΔACIAD1551::P<sub>tac</sub>-*lacZ*, ΔmutS::Spec<sup>R</sup>, ΔcomM | lacZ introduced into a defunct transposase (neutral gene = ACIAD1551), mutS deleted and replaced with Spec<sup>R</sup> (ACIAD1500), comM in-frame mutation (ACIAD0242) | This Study (TND0149/ SAD1326) |
| Δ180 (ADP1) | ΔmutS::Kan<sup>R</sup> | MutS deleted and replaced with a Kan<sup>R</sup> cassette | This Study (SAD742) |
| Δ180 ΔcomM (ADP1) | ΔmutS::Kan<sup>R</sup>, ΔcomM | MutS deleted and replaced with a Kan<sup>R</sup> cassette, ΔcomM in-frame | This Study (TND0144/ SAD1327) |
| Point mutant (ADP1) | ACIAD1551::Spec<sup>R</sup> (point mutant), ΔmutS::Kan<sup>R</sup> | SpecR cassette in ACIAD1551 is inactivated with a point mutation. MutS deleted and replaced with Kan<sup>R</sup> | This Study (TND0150/ SAD1328) |
| Point mutant, ΔcomM (ADP1) | ACIAD1551::Spec<sup>R</sup> (point mutant), ΔmutS::Kan<sup>R</sup>, ΔcomM | SpecR cassette in ACIAD1551 is inactivated with a point mutation. mutS was deleted and replaced with Kan<sup>R</sup> and comM was deleted in-frame | This Study (TND0164/ SAD1329) |
| pBAD18 Kan | pBAD18 Kan | TG1 E. coli strain used to purify a replicating plasmid with a Kan<sup>R</sup> cassette | This Study (SAD233) |
| P<sub>tac</sub>-*tfoX*, ΔradA | VC1153 OE Kan<sup>R</sup>, *radA*::Spec<sup>R</sup> | *radA* (VC2343) deleted and replaced with Spec<sup>R</sup> in a TfoX overexpressing background. | This Study (TMN0135/ SAD1813) |
| P<sub>tac</sub>-*tfoX*, ΔradA, ΔcomM | VC1153 OE Kan<sup>R</sup>, *radA*::Spec<sup>R</sup>, *comM*::Carb<sup>R</sup> | *comM* (VC0032) deleted and replaced with Carb<sup>R</sup> in a TfoX overexpressing, *radA* deletion background. | This Study (TMN0136/ SAD1814) |
| **Strains used in ComM Induction** |  |  |  |
| P<sub>tac</sub>-*gfp* | P<sub>tac</sub>-GFPmut3 Spec<sup>R</sup> at the *lacZ* locus | Replaced lacZ gene in SAD030 with a fragment from the transposon vector pDL1098 which encodes LacI, SpecR and has a P<sub>tac</sub> promoter. Cloned GFPmut3 downstream of P<sub>tac</sub> promoter | This Study (SAD559) |
| P<sub>tac</sub>-*gfp-comM* | P<sub>tac</sub> N-terminal *gfp-comM* at *lacZ* Spec<sup>R</sup>, ΔcomM Kan | VC0032 deletion strain containing an N-terminally GFP tagged comM at the *lacZ* locus under the control of P<sub>tac</sub> | This Study (SAD921) |
| *gfp-comM* | N-terminal GFP-comM at the native locus | N-terminally GFP tagged comM was cloned into SAD030 at the native locus | This Study (SAD924) |
| P<sub>tac</sub>-*tfoX*, *gfp-comM* | VC1153 OE Kan<sup>R</sup>, N-terminally GFP-comM at the native locus | Amplified VC1153 OE Kan from SAD061 and cloned fragment into SAD924 | This Study (TMN0140 / SAD1320) |
| $P_{tac-tfoX}$,<br>*gfp-comM<sup>K224A</sup>* | VC1153 OE *Kan<sup>R</sup>*, N-terminal *gfp-comM<sup>K224A</sup>* at the native locus, *Spec<sup>R</sup>* | Amplified *comM<sup>K224A</sup>* mutation from SAD1026 and cloned fragment into SAD1320 | This Study (TMN0148 / SAD1545) |
| **Strains Used for Protein Purification** |  |  |  |
| ComM | ComM cloned into *Amp<sup>R</sup>* StrepII expression vector | ComM N terminally tagged with 4x StrepII | This Study (pMB486 / SAD1330) |
| ComM<sup>K224A</sup> | ComM<sup>K224A</sup> cloned into *Amp<sup>R</sup>* StrepII expression vector | ComM<sup>K224A</sup> N terminally tagged with 4x StrepII | This Study (pMB488 / SAD1331) |
| Rosetta 2 (DE3) |  | *E. coli* expression strain | (pMB131/ SAD1332) |

**Table s2.**
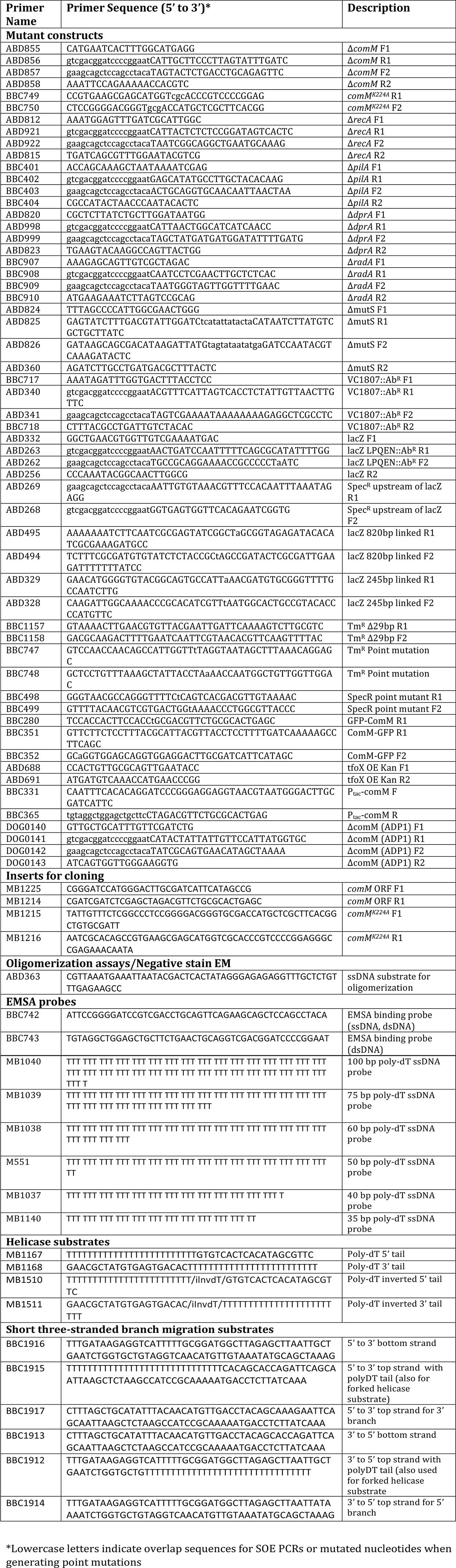
Primers used in this study.

## SUPPLEMENTARY METHODS

### Protein expression and purification

The *comM* open reading frame was PCR-amplified from *V. cholerae* genomic DNA using oligonucleotides MB1225 (CGGGATCCATGGGACTTGCGATCATTCATAGCCG) and MB1214 (CGATCGATCTCGAGCTAGACGTTCTGCGCACTGAGC), digested with *Bam*HI and *Xho*I, and ligated into the same sites in plasmid pMB131 to generate pMB486. This cloning added an N-terminal 4x Strep-tag II to the translated protein. The expression plasmid encoding the ATPase- and helicase-dead *comM-K224A* allele (pMB488) was created site-directed mutagenesis of pMB486 with oligonucleotides MB1215 (TATTGTTTCTCGGCCCTCCGGGGACGGGTGCGACCATGCTCGCTTCACGGCTGTGCGATT) and MB1216 (AATCGCACAGCCGTGAAGCGAGCATGGTCGCACCCGTCCCCGGAGGGCCGAGAAACAATA) and verified by DNA sequencing (ACGT, Inc.). Expression plasmids were transformed into Rosetta 2(DE3) pLysS cells and selected for at 37°C on LB medium supplemented with 100 μg/mL ampicillin and 34 μg/mL chloramphenicol. Fresh transformants were used to inoculate one or more 5-mL LB cultures supplemented with antibiotics and incubated at 30°C for ~6 h with aeration. These starter cultures were then diluted 1:100 in ZYP-5052 autoinduction medium containing 1x trace metals mix (3), 100 μg/mL ampicillin, and 34 μg/mL chloramphenicol and incubated at 22°C with agitation to OD600 >3 (15-18 h). Cells were harvested by centrifugation for 10 min at 5,500 x g and 4°C. Cell pellets were weighed and frozen at -80°C prior to lysis or for long-term storage.

Frozen cell pellets were thawed at room temperature by stirring in 4 mL/g cell pellet resuspension buffer (25 mM Na-HEPES (pH 7.5), 5% (v/v) glycerol, 300 mM NaOAc, 5 mM MgOAc, and 0.05% Tween-20) supplemented with 1x protease inhibitor cocktail (Sigma), and 20 μg/mL DNase I. Cells were lysed by six passed through a Cell Cracker operated at >1000 psi. All subsequent steps were performed at 4°C. The soluble fraction was clarified by centrifugation for 30 min at 33,000 x g followed by filtering the supernatant through a 0.22-μm membrane. This mixture was then applied to a Strep-Tactin Sepharose column (IBA) pre-equilibrated in resuspension buffer using an ÄKTA Pure (GE Healthcare Life Sciences). The column was washed with 20 column volumes (CVs) of resuspension buffer, 10 CVs of resuspension buffer supplemented with 5 mM ATP, and 10 CVs of resuspension buffer. Protein was eluted with 15 CVs of resuspension buffer supplemented with 2.5 mM desthiobiotin (IBA). Column fractions were examined on 8% SDS-PAGE gels run at 20 V/cm and stained with Coomassie Brilliant Blue R-250 (BioRad). Peak fractions were pooled, concentrated with Amicon Ultra-4 30K centrifugal filters, and loaded onto a HiPrep 16/60 Sephacryl S-200 HR column (GE Healthcare Life Sciences) pre-equilibrated in gel filtration buffer (25 mM Na-HEPES (pH 7.5), 5% glycerol, 300 mM NaCl, 5 mM MgCl2, and 0.05% Tween-20). The protein was eluted with 1.5 CVs gel filtration buffer, and fractions were analyzed by SDS-PAGE as above. Peak fractions were pooled, snap-frozen with liquid nitrogen, and stored at -80°C.

The *Saccharomyces cerevisiae* Pif1 helicase was overexpressed In Rosetta cells from plasmid pMB330 as described for ComM above. Pif1 purification was likewise identical, except the protein from the Strep-Tactin column was polished by Ni-affinity chromatography instead of size exclusion. Briefly, the pooled peak fractions were applied to a His60 Ni Superflow (Clontech) gravity column, washed with 10 CVs resuspension buffer supplemented with 25 mM imidazole (pH 8), and eluted with 4.5 CVs of a step gradient of resuspension buffer containing 100 mM, 250, and 500 mM imidazole (pH 8). Peak fractions were pooled, buffer exchanged into storage buffer (4), snap-frozen with liquid nitrogen, and stored at -80°C. The *Mycobacterium smegmatis* SftH was purified exactly as previously described (5).

ComM preps were tested for nuclease activity by incubating a labeled ssDNA probe with 100 nM of each protein prep in resuspension buffer for 1 hour at 37°C. Samples were then deproteinated with 1X stop load buffer and separated by native PAGE to assess degradation of the probe. All ComM protein preps used lacked detectable nuclease activity.

### Blue native PAGE

Oligomerization of ComM protein was assayed using Blue Native PAGE electrophoresis. 2.5 μM purified ComM was incubated for 30 min at room temperature in reaction buffer [10 mM Tris-HCl pH7.5, 20mM KCl, 1mM DTT, 10% Glycerol] with 5 mM ATP and/or 5 nM ssDNA (oligo ABD363) where indicated. 1 μL 20x sample buffer [5% Coomassie G-250, 0.5 M aminocaproic acid pH 7] was added to each reaction and samples were run on 4-16% Native PAGE gels [gel buffer = 0.5 M aminocaproic acid pH 7.0, 0.05 M Bis-Tris pH 7.0]. The cathode buffer was composed of 50 mM Tricine, 15 mM Bis-Tris pH 7.0, 0.02% Coomassie G-250, while the anode buffer was composed of 50 mM Bis-Tris pH 7.0. Samples were run at 150 V for 30 min, then 200 V for 45 min.

### Negative stain electron microscopy

The nominal magnification for the images is 60,000x, which is equivalent to 1.8 Å per pixel at the final image. Initial image processing, particle boxing, and CTF determination were performed using EMAN2 (6). A phase-flipped particle dataset was then imported into Relion (7) for 2D classification. Classes showing noisy images were discarded at this stage. As we observed clear six-fold symmetry from the classes, the subsequent processing imposed C6 symmetry. The remaining “good” classes were used to generate the initial models using e2initialmodel.py. The 3D classification was carried out using the initial model that was low-pass filtered to 40 Å to eliminate the possible effect from the model bias. Three 3D classes were obtained; the highest population (46%) of the classes was subjected to further structure refinement in Relion. Approximately 32,958 particles were used to generate the final 3D reconstruction. The reported resolution is ~13.8 Å using gold-standard Fourier shell correlation at a 0.143 cutoff; however, it is an over-estimated value because of the use of negative stain. The structure is rendered using UCSF Chimera (8).

### Helicase Assays

Fork substrates for helicase assays were made by 5’-end labelling oligonucleotides (**Table S2**) with T4 polynucleotide kinase (T4 PNK; NEB) and γ[32P]-ATP. Labelled oligonucleotides were separated from free label using illustra ProbeQuant G-50 micro columns (GE Healthcare) following the manufacturer’s instructions. Oligonucleotides were annealed by incubation with an equimolar amount of partially complementary oligonucleotides overnight at 37°C in annealing buffer (20 mM Tris-HCl [pH 8], 4% glycerol, 0.1 mM EDTA, 40 μg/mL BSA, 10 mM DTT, and 10 mM MgOAc) (9).

The DNA fork that allows for 5’-3’ and 3’-5’ activity was made by annealing oligonucelotides MB1167 with MB1168. The DNA fork that only allows for 5’-3’ helicase activity was made by annealing MB1167 / MB1511. The DNA fork that only allows for 3’-5’ helicase activity was made by annealing MB1168 / MB1510. DNA unwinding was assessed by incubating the indicated concentrations of helicase with 5 mM ATP and 0.1 nM radiolabelled fork in resuspension buffer. Reactions were incubated at 37°C for 30 min and stopped with the addition of 1x Stop-Load dye (5% glycerol, 20 mM EDTA, 0.05% SDS, and 0.25% bromophenol blue) supplemented with 400 μg/mL Proteinase K followed by a 10-min incubation at 37°C. Unwound DNA was then separated on 8% 19:1 acrylamide:bis-acrylamide gels in TBE buffer at 10 V/cm. Gels were dried under vacuum and imaged using a Typhoon 9210 Variable Mode Imager. DNA binding was quantified using ImageQuant 5.2 software.

## REFERENCES

1. Seitz, P. and Blokesch, M. (2013) Cues and regulatory pathways involved in natural competence and transformation in pathogenic and environmental Gram-negative bacteria. FEMS Microbiol Rev, 37, 336–363.

2. Meibom, K.L., Blokesch, M., Dolganov, N.A., Wu, C.Y. and Schoolnik, G.K. (2005) Chitin induces natural competence in Vibrio cholerae. Science, 310, 1824–1827.

3. Dalia, A.B., Lazinski, D.W. and Camilli, A. (2014) Identification of a membrane-bound transcriptional regulator that links chitin and natural competence in Vibrio cholerae. MBio, 5, e01028–01013.

4. Yamamoto, S., Mitobe, J., Ishikawa, T., Wai, S.N., Ohnishi, M., Watanabe, H. and Izumiya, H. (2014) Regulation of natural competence by the orphan two-component system sensor kinase ChiS involves a non-canonical transmembrane regulator in Vibrio cholerae. Mol Microbiol, 91, 326–347.

5. Lo Scrudato, M. and Blokesch, M. (2012) The regulatory network of natural competence and transformation of Vibrio cholerae. PLoS Genet, 8, e1002778.

6. Lo Scrudato, M. and Blokesch, M. (2013) A transcriptional regulator linking quorum sensing and chitin induction to render Vibrio cholerae naturally transformable. Nucleic Acids Res, 41, 3644–3658.

7. Metzgar, D., Bacher, J.M., Pezo, V., Reader, J., Doring, V., Schimmel, P., Marliere, P. and de Crecy-Lagard, V. (2004) Acinetobacter sp. ADP1: an ideal model organism for genetic analysis and genome engineering. Nucleic Acids Res, 32, 5780–5790.

8. Lorenz, M.G. and Wackernagel, W. (1994) Bacterial gene transfer by natural genetic transformation in the environment. Microbiol Rev, 58, 563–602.

9. Gwinn, M.L., Ramanathan, R., Smith, H.O. and Tomb, J.F. (1998) A new transformation-deficient mutant of Haemophilus influenzae Rd with normal DNA uptake. J Bacteriol, 180, 746–748.

10. Miller, V.L., DiRita, V.J. and Mekalanos, J.J. (1989) Identification of toxS, a regulatory gene whose product enhances toxR-mediated activation of the cholera toxin promoter. J Bacteriol, 171, 1288–1293.

11. Juni, E. and Janik, A. (1969) Transformation of Acinetobacter calco-aceticus (Bacterium anitratum). J Bacteriol, 98, 281–288.

12. Dalia, A.B., Lazinski, D.W. and Camilli, A. (2013) Characterization of undermethylated sites in Vibrio cholerae. J Bacteriol, 195, 2389–2399.

13. Dalia, A.B., McDonough, E. and Camilli, A. (2014) Multiplex genome editing by natural transformation. Proc Natl Acad Sci U S A, 111, 8937–8942.

14. Dalia, A.B. (2016) RpoS is required for natural transformation of Vibrio cholerae through regulation of chitinases. Environ Microbiol, 18, 3758–3767.

15. Studier, F.W. (2005) Protein production by auto-induction in high density shaking cultures. Protein Expr Purif, 41, 207–234.

16. Wittig, I., Braun, H.P. and Schagger, H. (2006) Blue native PAGE. Nat Protoc, 1, 418–428.

17. Wu, M. and Eisen, J.A. (2008) A simple, fast, and accurate method of phylogenomic inference. Genome Biol, 9, R151.

18. Edgar, R.C. (2004) MUSCLE: multiple sequence alignment with high accuracy and high throughput. Nucleic Acids Res, 32, 1792–1797.

19. Price, M.N., Dehal, P.S. and Arkin, A.P. (2010) FastTree 2--approximately maximum-likelihood trees for large alignments. PLoS One, 5, e9490.

20. Darling, A.E., Jospin, G., Lowe, E., Matsen, F.A.t., Bik, H.M. and Eisen, J.A. (2014) PhyloSift: phylogenetic analysis of genomes and metagenomes. PeerJ, 2, e243.

21. Stamatakis, A. (2014) RAxML version 8: a tool for phylogenetic analysis and post-analysis of large phylogenies. Bioinformatics, 30, 1312–1313.

22. Le, S.Q. and Gascuel, O. (2008) An improved general amino acid replacement matrix. Molecular biology and evolution, 25, 1307–1320.

23. Cooper, D.L. and Lovett, S.T. (2016) Recombinational branch migration by the RadA/Sms paralog of RecA in Escherichia coli. eLife, 5.

24. Mortier-Barriere, I., Velten, M., Dupaigne, P., Mirouze, N., Pietrement, O., McGovern, S., Fichant, G., Martin, B., Noirot, P., Le Cam, E. et al. (2007) A key presynaptic role in transformation for a widespread bacterial protein: DprA conveys incoming ssDNA to RecA. Cell, 130, 824–836.

25. Suckow, G., Seitz, P. and Blokesch, M. (2011) Quorum sensing contributes to natural transformation of Vibrio cholerae in a species-specific manner. J Bacteriol, 193, 4914–4924.

26. Hanson, P.I. and Whiteheart, S.W. (2005) AAA+ proteins: have engine, will work. Nat Rev Mol Cell Biol, 6, 519–529.

27. Kelley, L.A., Mezulis, S., Yates, C.M., Wass, M.N. and Sternberg, M.J. (2015) The Phyre2 web portal for protein modeling, prediction and analysis. Nat Protoc, 10, 845–858.

28. Brewster, A.S., Wang, G., Yu, X., Greenleaf, W.B., Carazo, J.M., Tjajadi, M., Klein, M.G. and Chen, X.S. (2008) Crystal structure of a near-full-length archaeal MCM: functional insights for an AAA+ hexameric helicase. Proc Natl Acad Sci U S A, 105, 20191–20196.

29. Kaplan, D.L., Davey, M.J. and O’Donnell, M. (2003) Mcm4,6,7 uses a “pump in ring” mechanism to unwind DNA by steric exclusion and actively translocate along a duplex. J Biol Chem, 278, 49171–49182.

30. Borgeaud, S., Metzger, L.C., Scrignari, T. and Blokesch, M. (2015) The type VI secretion system of Vibrio cholerae fosters horizontal gene transfer. Science, 347, 63–67.

31. Bochman, M.L. and Schwacha, A. (2009) The Mcm complex: unwinding the mechanism of a replicative helicase. Microbiol Mol Biol Rev, 73, 652–683.

32. Lloyd, R.G. and Sharples, G.J. (1993) Processing of recombination intermediates by the RecG and RuvAB proteins of Escherichia coli. Nucleic Acids Res, 21, 1719–1725.

33. West, S.C. (1997) Processing of recombination intermediates by the RuvABC proteins. Annu Rev Genet, 31, 213–244.

34. Paeschke, K., Bochman, M.L., Garcia, P.D., Cejka, P., Friedman, K.L., Kowalczykowski, S.C. and Zakian, V.A. (2013) Pif1 family helicases suppress genome instability at G-quadruplex motifs. Nature, 497, 458–462.

35. Kaplan, D.L. and O’Donnell, M. (2002) DnaB drives DNA branch migration and dislodges proteins while encircling two DNA strands. Mol Cell, 10, 647–657.

36. Yakovleva, L. and Shuman, S. (2012) Mycobacterium smegmatis SftH exemplifies a distinctive clade of superfamily II DNA-dependent ATPases with 3’ to 5’ translocase and helicase activities. Nucleic Acids Res, 40, 7465–7475.

37. Fallesen, T., Roostalu, J., Duellberg, C., Pruessner, G. and Surrey, T. (2017) Ensembles of Bidirectional Kinesin Cin8 Produce Additive Forces in Both Directions of Movement. Biophys J, 113, 2055–2067.

38. Marie, L., Rapisarda, C., Morales, V., Berge, M., Perry, T., Soulet, A.L., Gruget, C., Remaut, H., Fronzes, R. and Polard, P. (2017) Bacterial RadA is a DnaB-type helicase interacting with RecA to promote bidirectional D-loop extension. Nat Commun, 8, 15638.

39. Rozen, F., Edery, I., Meerovitch, K., Dever, T.E., Merrick, W.C. and Sonenberg, N. (1990) Bidirectional RNA helicase activity of eucaryotic translation initiation factors 4A and 4F. Mol Cell Biol, 10, 1134–1144.

40. Bugreev, D.V., Brosh, R.M., Jr. and Mazin, A.V. (2008) RECQ1 possesses DNA branch migration activity. J Biol Chem, 283, 20231–20242.

41. Whitby, M.C. and Lloyd, R.G. (1995) Branch migration of three-strand recombination intermediates by RecG, a possible pathway for securing exchanges initiated by 3’-tailed duplex DNA. EMBO J, 14, 3302–3310.

42. Burghout, P., Bootsma, H.J., Kloosterman, T.G., Bijlsma, J.J., de Jongh, C.E., Kuipers, O.P. and Hermans, P.W. (2007) Search for genes essential for pneumococcal transformation: the RADA DNA repair protein plays a role in genomic recombination of donor DNA. J Bacteriol, 189, 6540–6550.

43. Carrasco, B., Fernandez, S., Asai, K., Ogasawara, N. and Alonso, J.C. (2002) Effect of the recU suppressors sms and subA on DNA repair and homologous recombination in Bacillus subtilis. Mol Genet Genomics, 266, 899–906.

44. Weng, M.W., Zheng, Y., Jasti, V.P., Champeil, E., Tomasz, M., Wang, Y., Basu, A.K. and Tang, M.S. (2010) Repair of mitomycin C mono- and interstrand cross-linked DNA adducts by UvrABC: a new model. Nucleic Acids Res, 38, 6976–6984.

45. Lundin, C., North, M., Erixon, K., Walters, K., Jenssen, D., Goldman, A.S. and Helleday, T. (2005) Methyl methanesulfonate (MMS) produces heat-labile DNA damage but no detectable in vivo DNA double-strand breaks. Nucleic Acids Res, 33, 3799–3811.

46. Goranov, A.I., Kuester-Schoeck, E., Wang, J.D. and Grossman, A.D. (2006) Characterization of the global transcriptional responses to different types of DNA damage and disruption of replication in Bacillus subtilis. J Bacteriol, 188, 5595–5605.

47. Better, M. and Helinski, D.R. (1983) Isolation and characterization of the recA gene of Rhizobium meliloti. J Bacteriol, 155, 311–316.

48. Michod, R.E., Wojciechowski, M.F. and Hoelzer, M.A. (1988) DNA repair and the evolution of transformation in the bacterium Bacillus subtilis. Genetics, 118, 31–39.

49. Redfield, R.J. (1993) Genes for breakfast: the have-your-cake-and-eat-it-too of bacterial transformation. J Hered, 84, 400–404.

50. San Martin, M.C., Stamford, N.P., Dammerova, N., Dixon, N.E. and Carazo, J.M. (1995) A structural model for the Escherichia coli DnaB helicase based on electron microscopy data. J Struct Biol, 114, 167–176.

51. Gai, D., Zhao, R., Li, D., Finkielstein, C.V. and Chen, X.S. (2004) Mechanisms of conformational change for a replicative hexameric helicase of SV40 large tumor antigen. Cell, 119, 47–60.

52. Li, D., Zhao, R., Lilyestrom, W., Gai, D., Zhang, R., DeCaprio, J.A., Fanning, E., Jochimiak, A., Szakonyi, G. and Chen, X.S. (2003) Structure of the replicative helicase of the oncoprotein SV40 large tumour antigen. Nature, 423, 512–518.

53. Costa, A., Pape, T., van Heel, M., Brick, P., Patwardhan, A. and Onesti, S. (2006) Structural basis of the Methanothermobacter thermautotrophicus MCM helicase activity. Nucleic Acids Res, 34, 5829–5838.

54. Strycharska, M.S., Arias-Palomo, E., Lyubimov, A.Y., Erzberger, J.P., O’Shea, V.L., Bustamante, C.J. and Berger, J.M. (2013) Nucleotide and partner-protein control of bacterial replicative helicase structure and function. Mol Cell, 52, 844–854.

55. Shigenobu, S., Watanabe, H., Hattori, M., Sakaki, Y. and Ishikawa, H. (2000) Genome sequence of the endocellular bacterial symbiont of aphids Buchnera sp. APS. Nature, 407, 81–86.

56. Stephens, R.S., Kalman, S., Lammel, C., Fan, J., Marathe, R., Aravind, L., Mitchell, W., Olinger, L., Tatusov, R.L., Zhao, Q. et al. (1998) Genome sequence of an obligate intracellular pathogen of humans: Chlamydia trachomatis. Science, 282, 754–759.

57. Rocap, G., Larimer, F.W., Lamerdin, J., Malfatti, S., Chain, P., Ahlgren, N.A., Arellano, A., Coleman, M., Hauser, L., Hess, W.R. et al. (2003) Genome divergence in two Prochlorococcus ecotypes reflects oceanic niche differentiation. Nature, 424, 1042–1047.

58. Smith, M.G., Gianoulis, T.A., Pukatzki, S., Mekalanos, J.J., Ornston, L.N., Gerstein, M. and Snyder, M. (2007) New insights into Acinetobacter baumannii pathogenesis revealed by high-density pyrosequencing and transposon mutagenesis. Genes Dev, 21, 601–614.

59. Ramirez, M.S., Vilacoba, E., Stietz, M.S., Merkier, A.K., Jeric, P., Limansky, A.S., Marquez, C., Bello, H., Catalano, M. and Centron, D. (2013) Spreading of AbaR-type genomic islands in multidrug resistance Acinetobacter baumannii strains belonging to different clonal complexes. Curr Microbiol, 67, 9–14.

60. Traglia, G.M., Chua, K., Centron, D., Tolmasky, M.E. and Ramirez, M.S. (2014) Whole-genome sequence analysis of the naturally competent Acinetobacter baumannii clinical isolate A118. Genome Biol Evol, 6, 2235–2239.

61. Dalia, A.B., Seed, K.D., Calderwood, S.B. and Camilli, A. (2015) A globally distributed mobile genetic element inhibits natural transformation of Vibrio cholerae. Proc Natl Acad Sci U S A, 112, 10485–10490.

62. Konkol, M.A., Blair, K.M. and Kearns, D.B. (2013) Plasmid-encoded ComI inhibits competence in the ancestral 3610 strain of Bacillus subtilis. J Bacteriol, 195, 4085–4093.

63. Rabinovich, L., Sigal, N., Borovok, I., Nir-Paz, R. and Herskovits, A.A. (2012) Prophage excision activates Listeria competence genes that promote phagosomal escape and virulence. Cell, 150, 792–802.

64. Croucher, N.J., Harris, S.R., Barquist, L., Parkhill, J. and Bentley, S.D. (2012) A high-resolution view of genome-wide pneumococcal transformation. PLoS Pathog, 8, e1002745.

65. Mell, J.C., Shumilina, S., Hall, I.M. and Redfield, R.J. (2011) Transformation of natural genetic variation into Haemophilus influenzae genomes. PLoS Pathog, 7, e1002151.

66. Al-Deib, A.A., Mahdi, A.A. and Lloyd, R.G. (1996) Modulation of recombination and DNA repair by the RecG and PriA helicases of Escherichia coli K-12. J Bacteriol, 178, 6782–6789.

67. Kline, K.A. and Seifert, H.S. (2005) Mutation of the priA gene of Neisseria gonorrhoeae affects DNA transformation and DNA repair. J Bacteriol, 187, 5347–5355.

68. Kruger, N.J. and Stingl, K. (2011) Two steps away from novelty-principles of bacterial DNA uptake. Mol Microbiol, 80, 860–867.

69. Martin, B., Sharples, G.J., Humbert, O., Lloyd, R.G. and Claverys, J.P. (1996) The mmsA locus of Streptococcus pneumoniae encodes a RecG-like protein involved in DNA repair and in three-strand recombination. Mol Microbiol, 19, 1035–1045.

70. Whitby, M.C., Ryder, L. and Lloyd, R.G. (1993) Reverse branch migration of Holliday junctions by RecG protein: a new mechanism for resolution of intermediates in recombination and DNA repair. Cell, 75, 341–350.

71. Azeroglu, B., Mawer, J.S., Cockram, C.A., White, M.A., Hasan, A.M., Filatenkova, M. and Leach, D.R. (2016) RecG Directs DNA Synthesis during Double-Strand Break Repair. PLoS Genet, 12, e1005799.

72. Cox, M.M., Morrical, S.W. and Neuendorf, S.K. (1984) Unidirectional branch migration promoted by nucleoprotein filaments of RecA protein and DNA. Cold Spring Harb Symp Quant Biol, 49, 525–533.

73. Cox, M.M. and Lehman, I.R. (1981) Directionality and polarity in recA protein-promoted branch migration. Proc Natl Acad Sci U S A, 78, 6018–6022.

74. Papenbrock, J., Mock, H.P., Tanaka, R., Kruse, E. and Grimm, B. (2000) Role of magnesium chelatase activity in the early steps of the tetrapyrrole biosynthetic pathway. Plant Physiol, 122, 1161–1169.

## SI REFERENCES

1. Miller, V.L., DiRita, V.J. and Mekalanos, J.J. (1989) Identification of toxS, a regulatory gene whose product enhances toxR-mediated activation of the cholera toxin promoter. J Bacteriol, 171, 1288–1293.

2. Dalia, A.B., Lazinski, D.W. and Camilli, A. (2014) Identification of a membrane-bound transcriptional regulator that links chitin and natural competence in Vibrio cholerae. MBio, 5, e01028–01013.

3. Studier, F.W. (2005) Protein production by auto-induction in high density shaking cultures. Protein Expr Purif, 41, 207–234.

4. Paeschke, K., Bochman, M.L., Garcia, P.D., Cejka, P., Friedman, K.L., Kowalczykowski, S.C. and Zakian, V.A. (2013) Pif1 family helicases suppress genome instability at G-quadruplex motifs. Nature, 497, 458–462.

5. Yakovleva, L. and Shuman, S. (2012) Mycobacterium smegmatis SftH exemplifies a distinctive clade of superfamily II DNA-dependent ATPases with 3’ to 5’ translocase and helicase activities. Nucleic Acids Res, 40, 7465–7475.

6. Tang, G., Peng, L., Baldwin, P.R., Mann, D.S., Jiang, W., Rees, I. and Ludtke, S.J. (2007) EMAN2: an extensible image processing suite for electron microscopy. Journal of structural biology, 157, 38–46.

7. Scheres, S.H. (2012) RELION: implementation of a Bayesian approach to cryo-EM structure determination. Journal of structural biology, 180, 519–530.

8. Pettersen, E.F., Goddard, T.D., Huang, C.C., Couch, G.S., Greenblatt, D.M., Meng, E.C. and Ferrin, T.E. (2004) UCSF Chimera-a visualization system for exploratory research and analysis. J Comput Chem, 25, 1605–1612.

9. Kanter, D.M. and Kaplan, D.L. (2011) Sld2 binds to origin single-stranded DNA and stimulates DNA annealing. Nucleic Acids Res, 39, 2580–2592.

